# Rational design of a foldon-derived heterotrimer guided by quantitative native mass spectrometry

**DOI:** 10.1101/2025.03.25.645370

**Authors:** Xinyu Liu, David S. Roberts, Craig A. Bingman, Song Jin, Ying Ge, Samuel H. Gellman

## Abstract

Designing stable hetero-oligomeric protein complexes with defined inter-subunit stoichiometries remains a significant challenge. In this study, we report the design of a highly selective heterotrimeric assembly derived from the well-known foldon homotrimer. We generated an *aab* heterotrimer by introducing the Q11E modification to destabilize the homotrimer and a compensatory modification, either V14A or V14L, to stabilize the hydrophobic core of the heterotrimer. Native mass spectrometry (MS) was essential for guiding the design process, enabling precise characterization of oligomeric states and their equilibrium distributions. The heterotrimer structure was validated by x-ray crystallography. Our findings highlight the effectiveness of combining rational design with native MS to develop specific hetero-oligomeric assemblies.

## INTRODUCTION

Many cellular processes rely on the precise assembly of protein subunits into specific quaternary structures that drive biological function.^1^ Designing novel protein structural motifs that assemble into stable and well-defined oligomers can expand the toolkit for protein engineering.^2–5^ Considerable progress has been made in the development of protein homo-oligomers with defined stoichiometry, but hetero-oligomer design has proven significantly more challenging.^6–9^

Designed heterotrimers have so far been limited to two motifs, coiled coils and collagen-like triple helices.^9–13^ New approaches that harness alternative structural motifs and allow distinctive spatial arrangements of appended protein modules are needed to expand heterotrimer design. Here we describe first steps toward that goal based on the foldon unit, which is derived from the 27-residue C-terminal domain of bacteriophage T4 fibritin. This diminutive motif forms a very stable homotrimer that anchors 400 Å filaments of the bacteriophage.^14,15^ The native foldon trimer has been leveraged for the design of vaccines and biomaterials.^16–19^ Foldon derivatives that formed discrete and predictable heterotrimers would allow formation of new types of defined assemblies that bring different appended protein modules together.

We envisioned that carefully designed foldon variants could maintain a native-like tertiary structure but favor heterotrimer over homotrimer quaternary structure. The native foldon unit is highly sensitive to sequence perturbations, and all reported sequence modifications have destabilized the homotrimer assembly.^20^ The challenge we perceived was to identify a pair of variants with low homotrimerization propensity that could co-assemble to form a stable heterotrimer.

Biophysical techniques commonly employed to elucidate protein assembly behavior, such as X-ray crystallography, cryo-electron microscopy (cryo-EM), analytical ultracentrifugation, or nuclear magnetic resonance (NMR) spectroscopy, are time-consuming and therefore not ideal for guiding an iterative design process. We envisioned that native mass spectrometry (MS) would support rapid and incisive analysis of foldon variant assembly behavior.^21^ Previous work has shown that under appropriate nondenaturing conditions, information on protein complex composition, subunit stoichiometry, inter-subunit connectivity, and assembly dynamics obtained by a native MS experiment correlate with structural data available from cryo-EM, X-ray crystallography or NMR.^22–29^

Here we show how rational design guided by native MS enabled the discovery of a pair of foldon sequence variants that preferentially form an *aab* heterotrimer. The structure of this heterotrimer was established by X-ray crystallography.

## RESULTS AND DISCUSSION

### Experimental Design

In solution, the native foldon 27-mer (**Figure 1**) is known to display a dynamic equilibrium^15^ between homotrimers (*aaa*) and monomers (*a*), with the homotrimer strongly favored. We found that this equilibrium could be readily characterized via native MS (**Figure 1D-F**). The data revealed isotopically resolved monomers (*z* = 1+), homodimers (*z* = 3+ and 4+) and homotrimers (*z* = 4+ to 6+), with a strong bias toward homotrimer formation (**Figure 1E**). Because of the redundancy of isotopologues when accounting for multiple subunits of the same chemical formula, the different oligomeric states (*i*.*e*. monomer, homodimer, and homotrimer) can sometimes overlap at specific charge states, such as the case with the z = 6+ and z = 4+ charge states for the foldon homotrimer and homodimer, respectively. However, in all cases, the contribution of individual oligomeric states to a given experimental spectrum can be deconvoluted to obtain a global understanding of the oligomeric landscape.^30–32^ Deconvoluted MS of all isotopically resolved charge states yielded highly accurate masses (< 1 ppm mass error) commensurate with the expected foldon oligomeric states: monomer (3081.444 Da), homodimer (6162.890 Da), and homotrimer (9244.334 Da). Quantification of the abundance of each species revealed that the homotrimer was dominant (∼ 93 %), with only minor contributions by monomer (∼ 3%) or homodimer (∼4 %) species (**Figure 1F**). This analysis highlights the ability of native MS to profile the global oligomeric landscape and quantitatively define the oligomer populations.

**Figure 1.**
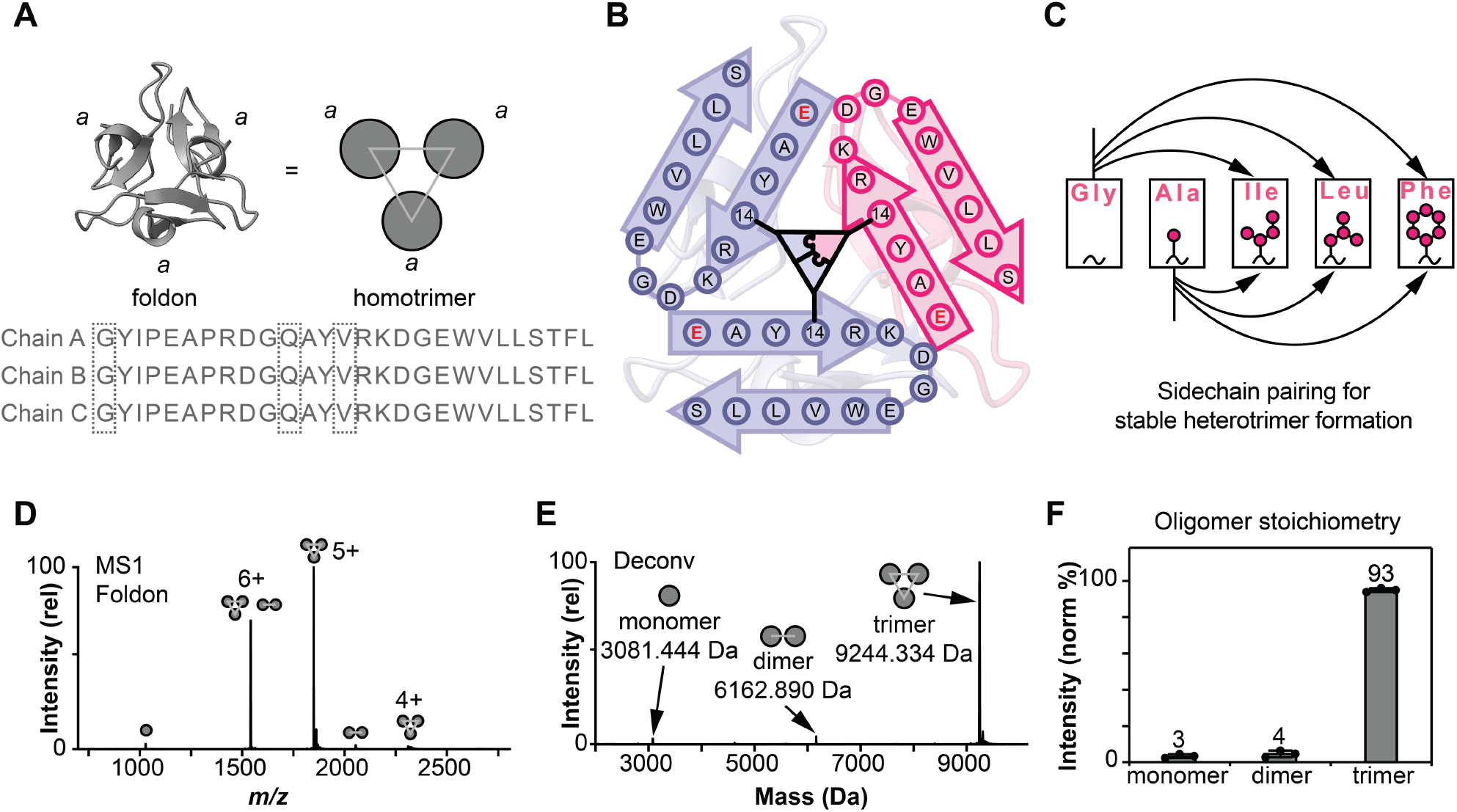
Designing a minimal heterotrimeric β-hairpin-propeller-like scaffold guided by native MS. **A.**Structure and amino acid sequence of foldon. The native structure of foldon consists of a stable homotrimeric scaffold (*aaa*) in solution. **B.** Biased formation of an *aab* heterotrimer can be achieved by introducing an optimal-sized hydrophobic packing at the oligomer interface. We hypothesized that the highlighted mutation Q11E will lead to partial trimer dissociation without ablation of the overall tertiary structure. **C.** Hydrophobic amino acids with complementary volumes classes at position 14 are potential to promote a biased packing pattern. Based on the ImMunoGeneTics information system,^40^ we screened all “small” and “large” volume class pairs at position 14 by generating the designer mutants to obtain a biased heterotrimer system. **D.** Native MS1 of Foldon. The 5+ charge state for the trimer peptide samples under native MS provides an isolated in m/z space such that they avoid any contributions from overlapping oligomeric states, which allows for straightforward identification of trimers with different mass. **E.** The native deconvoluted MS provides a direct way to analyze the relative abundance of various oligomeric states. **F.** The oligomer stoichiometry was quantified by deconvoluted MS. As expected, foldon exists dominantly as a trimer in solution. Data are representative of n = 3 independent experiments with error bars indicating the standard deviation.

We evaluated two targeted changes to the native foldon sequence before attempting to design a pair of variants that would preferentially form a heterotrimer. First, we replaced native residue Gly1 with Lys in order to enhance solubility without destabilizing the homotrimer. High-resolution structural data suggest that Gly1 does not contribute to homotrimer stability. As expected, native MS analysis revealed that the G1K modification did not significantly alter trimer stability (**Figure S1**).

The second change was replacement of native Gln11 with Glu. We predicted that this change would diminish homotrimer stability because in the native homotrimer, Gln11 on one protomer lies near Asp17 of another. Thus, Q11E should introduce Coulombic repulsion at the trimer interface. The foldon variant containing both G1K and Q11E formed a homotrimer that was less stable than the native foldon homotrimer, as indicated by native MS (lower proportion of homotrimer; **Figure 2A and Figure S2A**). Native foldon, the G1K variant, and the G1K-Q11E variant displayed very similar far-UV circular dichroism (CD) signatures, which suggests that the have very similar secondary structure (**Figure S3A**). Variable-temperature CD analysis showed that the native foldon and G1K variant were indistinguishable in terms of conformational stability (T_m_ = 348 K for both), but that the G1K-Q11E variant was less conformationally stable (T_m_ = 337 K) (**Figure S3B and Table S2**).

**Figure 2.**
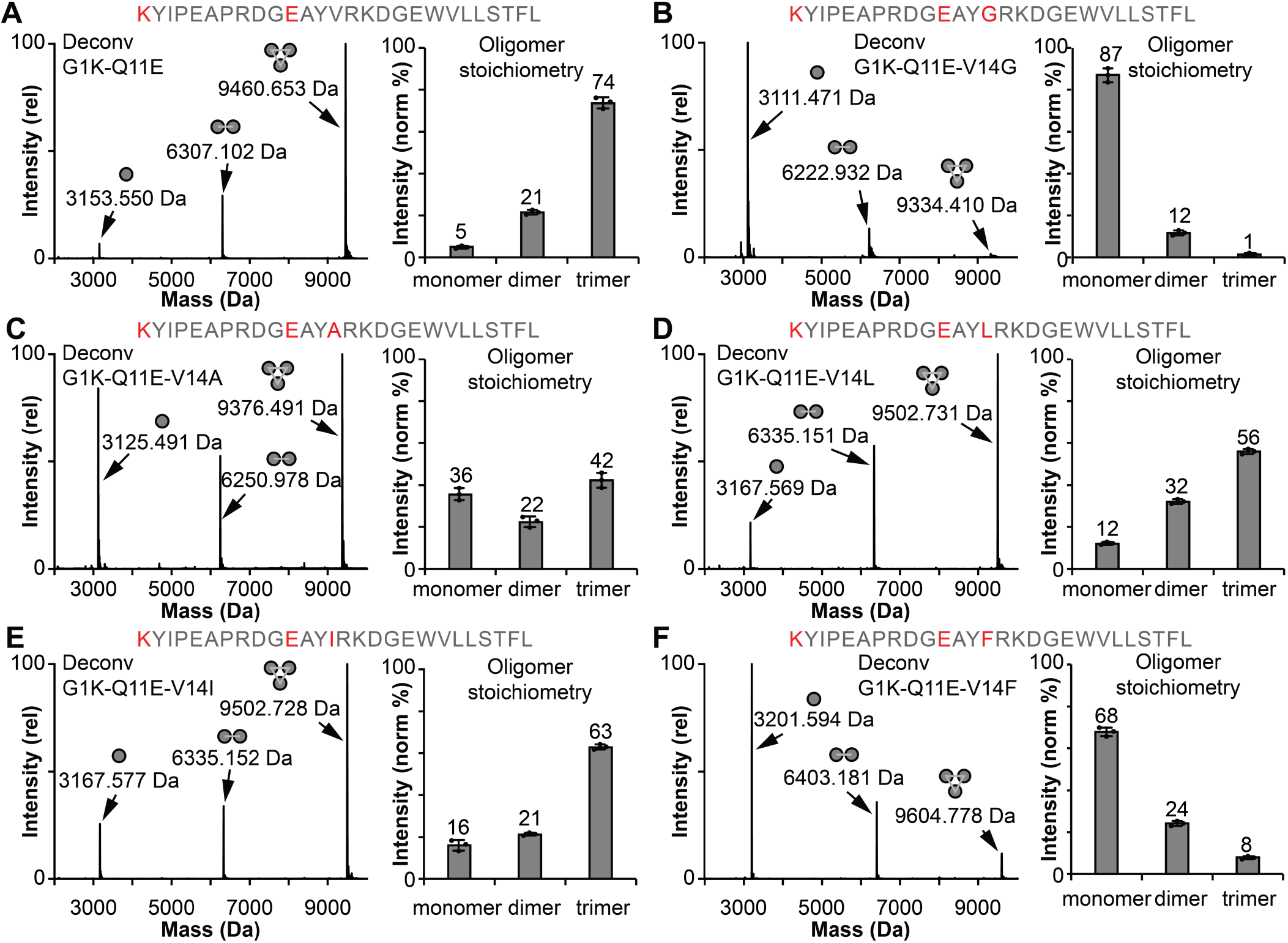
Native deconvoluted MS characterization of six peptides on derived from the foldon sequence. The corresponding MS1 spectra are included in **Figure S3.**The quantifications of oligomers are presented as bar graphs. **A.** G1K-Q11E shows a partially dissociated quaternary structure compared to the native foldon while maintaining a preferred trimeric structure. **B-F.** Variants containing the G1K and Q11E modifications along with a change at position 14, which forms the trimer core with the native Val, display different oligomer distributions relative to the native foldon. As predicted, residues with side chains that are very small (Gly) or large (Phe) form little trimer. On the other hand, Ala, Leu, and Ile each maintained a preferred trimeric state, but at a lower proportion relative to the native foldon.

High-resolution structures of the native foldon trimer reveal that the interface among the three protomers is dominated by the segment Q11-D17 (**Figure 1B**). The side chains of the three V14 residues pack intimately at the center of this interface. It seemed likely that foldon variants with larger or smaller side chains at position 14 might be hindered from self-assembly. We hypothesized that a pair of such variants might be able to form an *aab* heterotrimer in which two smaller side chains could be complemented at the trimer core by a single larger side chain (**Figure 1B-C**). This hypothesis was tested by examining a set of triple variants of the foldon sequence.

Each contained the G1K and Q11E modifications along with a modification at position 14 that varied the size of the nonpolar side chain. Within this set of triple variants, V14 was replaced by Gly, Ala, Ile, Leu or Phe.

### Foldon Triple Variants with variable propensities for homotrimer formation

The five new foldon variants, G1K-Q11E-V14G, G1K-Q11E-V14A, G1K-Q11E-V14L, G1K-Q11E-V14I, and G1K-Q11E-V14F, were each initially evaluated in isolation via native MS (**Figure 2B-F and Figure S2B-F)**. The variants possessing either the smallest (Gly) or largest (Phe) side chain at position 14 displayed nearly complete homotrimer dissociation, with a total relative homotrimer abundance of 1 % or 8 %, respectively (**Figure 2B and 2F)**. For the V14G variant, homotrimer destabilization may result from loss inter-subunit hydrophobic packing, or enhanced conformational flexibility that results from the Val→Gly replacement, or both factors. For the V14F variant, it seems likely that steric repulsions among the Phe side chains would prevent adoption of a native-like trimer quaternary structure.

Each of the remaining three foldon variants, G1K-Q11E-V14A, G1K-Q11E-V14L, and G1K-Q11E-V14I, displayed a substantial homotrimer population (42%-63%), but these populations were diminished relative to the homotrimer populations for native foldon, the G1K variant or the G1K-Q11E variant. Thus, the homotrimer formed by each of these three triple variants is less stable than the native foldon homotrimer.

### Pairing of Foldon Triple Variants in Search of a Stable Heterotrimer

In a system at equilibrium containing equal concentrations of foldon variants *a* and *b* that have comparable propensities for homotrimer and heterotrimer formation, we would expect uniform distribution of possible heterotrimer states to result in a 1:3:3:1 (*aaa*:*aab*:*abb*:*bbb*) mixture of trimers (**Figure 3A**).^33^ We tested this hypothesis using a 1:1 mixture of native foldon and the G1K variant. As anticipated, native MS analysis of this mixture revealed the presence of all four possible heterotrimer states (**Figure 3B**). We note that for all heterotrimer analysis, the most abundant trimeric species charge state (*z* = 5+) is sufficiently isolated in *m/z* that contributions from other oligomeric states are avoided; thus, it is straightforward to determine the composition of each trimer from data for this charge state.

**Figure 3.**
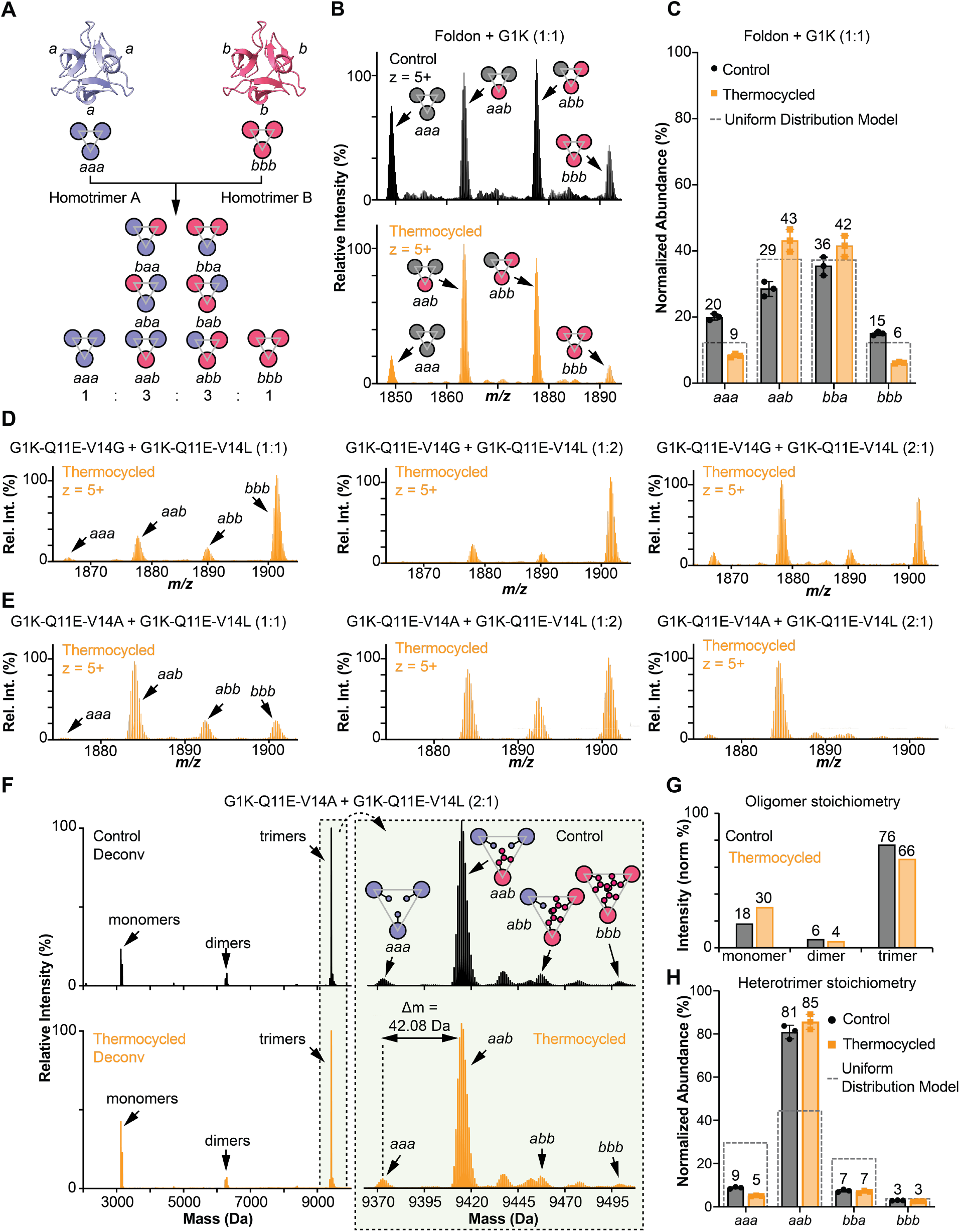
Identification and characterization of an *aab* hetero-oligomer. **A.** Mixing peptides with equal propensities to form homotrimers or heterotrimers generates four possible trimers with a 1:3:3:1 ratio, accounting for mass degeneracy. **B.** Zoomed-in native MS of 1:1 foldon (*a*) + G1K (*b*) to the charge state 5+ region showing the isotopic distributions of trimers with their assigned compositions (*aaa, aab*, etc.). The thermocycled spectrum shows a modified distribution, which presumably reflects the equilibrium state. **C.** Quantification of trimers compared to a uniform distribution model. The G1K modification does not affect the oligomerization of foldon. **D.** Zoomed-in native MS of 1:1, 1:2, and 2:1 G1K-Q11E-V14G (*a*) + G1K-Q11E-V14L (*b*) showing the isotopic distributions of trimers in the charge state 5+ region with their assigned compositions (*aaa, aab*, etc.). The thermocycled spectra show that G1K-Q11E-V14L stays dominantly as a homotrimer when mixed with G1K-Q11E-V14G. When the relative abundance of G1K-Q11E-V14G increases, the *aab* heterotrimer population increases; however, significant *bbb* homotrimer remains. **E.** Zoomed-in native MS of 1:1, 1:2, and 2:1 G1K-Q11E-V14A (*a*) + G1K-Q11E-V14L (*b*) to the charge state 5+ region showing the isotopic distributions of trimers with their assigned compositions (*aaa, aab*, etc.). The thermocycled spectra show that the *aab* heterotrimer is preferred when the monomers *a* and *b* have 2:1stoichiometry. **F.** Native deconvoluted MS of 2:1 G1K-Q11E-V14A (*a*) + G1K-Q11E-V14L (*b*) suggested a dominant population of trimers. The zoomed-in spectrum to the trimer region showed a dominant *aab* heterotrimer. The mass difference between each trimer signal, 42.08 Da, matches with the theoretical mass difference between Ala and Leu, which is 42.05 Da. **G.** Oligomer proportions of 2:1 G1K-Q11E-V14A + G1K-Q11E-V14L. After thermocycling, some trimers were dissociated to monomers. **H.** Trimer proportions of 2:1 G1K-Q11E-V14A + G1K-Q11E-V14L. The relative distribution of trimers remained unchanged after heating, both suggesting a dominant *aab* heterotrimer population, compared to the uniform distribution model.

To determine whether the population profile of the four trimeric species formed upon mixing equimolar native foldon and G1K variant at room temperature reflects an equilibrium state, this mixture was heated to 95°C for 30 min and then cooled to 4°C before reanalysis (**Figure 3B and 3C**). Trimer proportions changed in a small but significant way after heating, which suggests that the initial mixture was close to equilibrium, but not fully equilibrated. After heating, the trimer population was 9%:43%:42%:6% (*aaa*:*aab*:*bba*:*bbb*), which is similar to the 12.5%:37.5%:37.5%:12.5% population predicted if there is no energetic difference among the four trimer species for a 1:1 mixture foldon variants (**Figure 3C**). The slight preference for heterotrimers over homotrimers might result from weak Coulombic repulsions in the G1K homotrimer.

In an effort to discover a combination of foldon variants that would favor a single heterotrimeric composition, we examined all six pairings of a small side chain at position 14 (Gly or Ala) with a large side chain at position 14 (Ile, Leu or Phe). Each binary mixture was evaluated in a 2:1, 1:1 or 1:2 proportion. The efficiency of the native MS approach facilitated this thorough survey. Trimer population profile data for these and other combinations are provided in the Supporting Information. Here we discuss just two of these systems to highlight the insights available from native MS data.

First, we evaluated the binary mixtures of G1K-Q11E-V14G and G1K-Q11E-V14L (**Figure 3D**). At all proportions, the V14L variant predominantly forms a homotrimer (*bbb*). Although the stability for this homotrimer is significantly lower than for the native foldon trimer, as shown in **Figure 1 and 2**, the G1K-Q11E-V14L is sufficiently stable to exclude V14G protomers in a 1:1 mixture. When the relative abundance of the V14G variant was increased to 2:1, we observed an increase of the *aab* heterotrimer. However, even in this case we did not observe a significant preference for *aab* over *bbb*.

A similar analysis was undertaken with G1K-Q11E-V14A + G1K-Q11E-V14L (**Figure 3E**). At all proportions, the incorporation of V14A disrupts the homotrimeric V14L packing. In the 2:1 mixture, a single heterotrimer state (*aab*) was dominant. Quantitative deconvoluted MS (**Figure 3F**) revealed that thermocycling the 2:1 mixture resulted in < 10% change in the trimer profile (**Figure 3G**). At equilibrium, > 80% of the total trimer population corresponded to *aab* (**Figure 3H**). For a 2:1 mixture of monomers *a* and *b*, a uniform distribution of trimers would display a 29.6%:44.4%:22.2%:3.7% distribution for *aaa*:*aab*:*abb*:*bbb*.^33^ The heterotrimer distribution established by native MS, 9%:81%:7%:3% for *aaa*:*aab*:*abb*:*bbb*, is quite different and clearly establishes the intrinsic preference for the *aab* heterotrimer assembly in this system.

Variable temperature CD analysis of the 2:1 G1K-Q11E-V14A + G1K-Q11E-V14L mixture revealed a T_m_ = 316 K, which lies between the T_m_ values of homotrimer G1K-Q11E-V14A (T_m_ = 314 K) and homotrimer G1K-Q11E-V14L (T_m_ = 318 K) (**Figure S12 and Table S2**). Despite the lower thermal stability of the 2:1 mixture compared to the parent foldon structure (T_m_ = 348 K), the 2:1 binary mixture leads to highly selective formation of a discrete heterotrimer.

To provide a molecular rationale for the observed structural bias favoring *aab* formation in the 2:1 G1K-Q11E-V14A + G1K-Q11E-V14L mixture, we performed additional studies to investigate: (1) the effect of replacing Gly1 with Lys, in the absence of the Q11E modification, and (2) the effect of replacing Gln11 with Glu in the absence of the G1K modification (**Figure S13**). Native MS revealed that, without the G1K modification, the 2:1 mixture of Q11E-V14A + Q11E-V14L remains strongly biased toward *aab* heterotrimer formation (∼ 85%), suggesting that the G1K modification does not significantly impact heterotrimer selectivity (**Figure S13A**). However, the absence of the Q11E modification, in the 2:1 mixture of G1K-V14A + G1K-V14L, resulted in substantial reduction of *aab* trimer formation (∼ 55%) (**Figure S13B**). Together these results suggest that the combination of modifications to the end-to-end contacts (*i*.*e*., the Q11E modification) and the central hydrophobic cluster (*i*.*e*., the V14A and V14L modifications) are crucial for promoting formation of a specific heterotrimer.

### Structural elucidation of *aab* heterotrimer by X-ray crystallography

We co-crystallized G1K-Q11E-V14A and G1K-Q11E-V14L from a 2:1 stoichiometric mixture to enable X-ray crystallography (**Figure 4A**). The crystal structure, obtained at 1 Å resolution (PDB: 8UDN), revealed a heterotrimeric (*aab*) β-propeller-like structure (*P*43 space group) (see **Table S3** for details). Superimposing the resulting *aab* crystal structure with the solution-phase NMR structure of the native foldon (PDB: 1RFO)^15^ revealed similar quaternary structures (**Figure S15**). X-ray diffraction from the *aab* crystal structure indicated a high degree of localized disorder at the center of the heterotrimer, near position 14 (**Figure 4A**). We hypothesize that the partially disordered core results from rotational distortion owing to the isomorphic orientations of the heterotrimeric assembly (*i*.*e*., *aab, aba*, and *baa*) within the crystal lattice. Indeed, when the side chains at position 14 in the *aab* crystal structure were depicted as overlapping Ala14 and Leu14, we observed well-fitted electron density maps that agreed with our original hypothesis of an optimal-sized packing arrangement (**Figure S16**).

**Figure 4.**
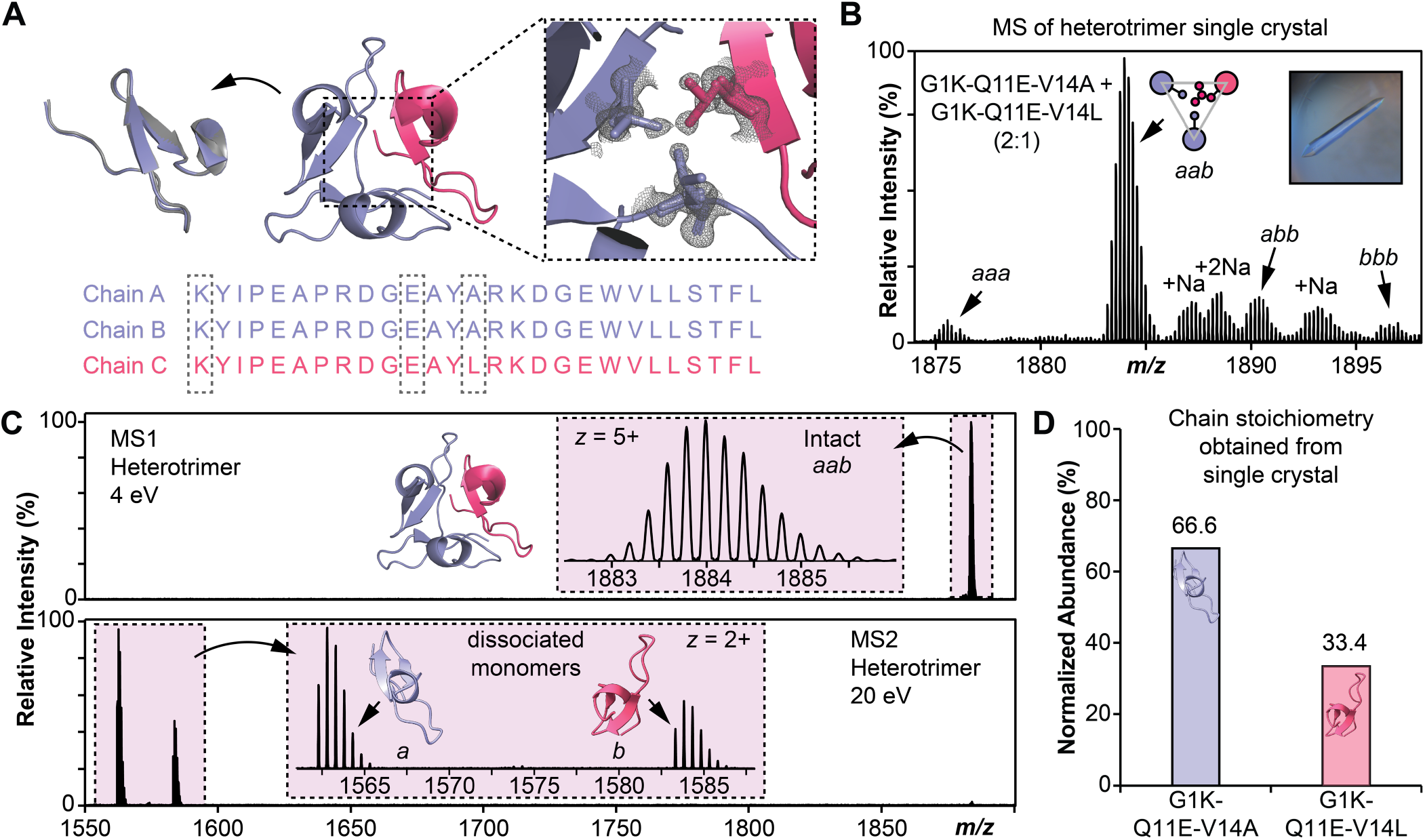
Structural elucidation of *aab* heterotrimer from 2:1 G1K-Q11E-V14A + G1K-Q11E-V14L by combined X-ray crystallography and native MS. **A.** Overlay of a protomer from the new heterotrimer PDB: 8UDN (blue) and a crystal structure of Foldon PDB: 4NCU^41^ (grey) shows a Root Mean Square Deviation (RMSD) of 0.305 Å, indicating similar overall structure of the homotrimer and heterotrimer. The electron density map at position 14 from the heterotrimer crystal. The high degree of localized disorder at the center of the heterotrimer structure compared to a homotrimer (**Figure S17**) aligns with the hypothesis that the crystal structure of a heterotrimer presents a disordered arrangement of *aab, aba*, and *baa* trimers within the crystal lattice. **B.** Zoomed-in native MS of solutions generated from crystals of 2:1 G1K-Q11E-V14A (*a*) + G1K-Q11E-V14L (*b*) suggested a dominant *aab* heterotrimer. The inset shows a representative image of the crystal arising from the 2:1 G1K-Q11E-V14A + G1K-Q11E-V14L mixture. **C.** Complex-up MS of the crystal-derived solution. Complex dissociation was achieved by applying a collision energy of 20 eV, yielding G1K-Q11E-V14A and G1K-Q11E-V14L monomers. **D.** The complex-up analysis revealed a 2:1 stoichiometry of the two monomers, which is consistent with the conclusion that the crystals obtained from the G1K-Q11E-V14A and G1K-Q11E-V14L 2:1 stoichiometric mixture contain the *aab* heterotrimer.

As shown above, the introduction of the Q11E modification resulted in partial dissociation of the trimer structure. The crystal structure does not indicate the inter-subunit repulsion we originally proposed between Glu11 and Asp17. However, we rationalize that the destabilizing effect of the Q11E modification might be a result of the close proximity between Asp9 and Glu11, which introduces intramolecular repulsion in the N-terminal loop segment, and thus can lead to distortion of the protomer conformation. Indeed, we observed an increase in the distance between side chains Asp9 and Glu11 in the *aab* heterotrimer (6.6 Å), as compared to the native foldon (3.7 Å) (**Figure S17**).

To assess the protomer stoichiometry within single crystals arising from the 2:1 mixture of G1K-Q11E-V14A and G1K-Q11E-V14L, we harvested multiple single crystals and dissolved them in 150 mM aqueous ammonium acetate. Native MS analysis of this solution revealed the *aab* state to be > 80% of the total trimer species (**Figure 4B**). This trimer state distribution is identical to the distribution observed after solution-phase equilibration (**Figure 3**). To gain further insight on the identity of the dominant heterotrimer this crystal-derived solution, we performed a complex-up^34^ analysis of the intact heterotrimer and characterized the dissociated subunits (**Figure 4C**). Following specific isolation of the heterotrimer species centered at 1884 *m/z* ± 2 *m/z*, complete complex dissociation was achieved by applying a collision energy of 20 eV, which yielded G1K-Q11E-V14A and G1K-Q11E-V14L monomers. The resulting stoichiometric abundance of the dissociated monomers was found to be a near perfect 2:1 ratio (66.6% G1K-Q11E-V14A and 33.4% G1K-Q11E-V14L) (**Figure 4D**). This result is consistent with our conclusion that the species crystallized was the *aab* heterotrimer comprised of two G1K-Q11E-V14A protomers and one G1K-Q11E-V14L protomer. The alignment between crystal structure and native MS analysis for the 2:1 mixture should motivate further utilization of quantitative native MS as a high-throughput and high-resolution technique for guiding the rational design of protein assemblies.^35–39^

## CONCLUSIONS

We have shown that the well-studied foldon homotrimer provides a basis for the design of sequence variants that form a specific heterotrimer. Prior efforts to design heterotrimers with specific stoichiometries have focused on two motifs, coiled coils and collage-like triple helices. Our findings suggest that foldon-derived systems will provide a fruitful alternative to these precedents that support new geometries among appended protein units. This study demonstrates the power of native MS as a technique for rapid and quantitative analysis of assemblies formed in solutions containing polypeptide mixtures.

## AUTHOR INFORMATION

### Authors

**Xinyu Liu —** Department of Chemistry, University of Wisconsin−Madison, Madison, Wisconsin 53706, United States;

**David S. Roberts —** Department of Chemistry, University of Wisconsin−Madison, Madison, Wisconsin 53706, United States;

**Craig A. Bingman —** Department of Biochemistry, University of Wisconsin−Madison, Madison, Wisconsin 53706, United States;

**Song Jin —** Department of Chemistry, University of Wisconsin−Madison, Madison, Wisconsin 53706, United States;

**Ying Ge —** Department of Chemistry, University of Wisconsin−Madison, Madison, Wisconsin 53706, United States; Department of Cell and Regenerative Biology, University of Wisconsin -Madison, Madison, Wisconsin 53705, United States; Human Proteomics Program, School of Medicine and Public Health, University of Wisconsin, Madison, Wisconsin 53705, United States;

### Present Addresses

X.L. — Department of Chemistry, Stanford University, Stanford, California 94305, United States

D.S.R. — Department of Chemistry and Sarafan ChEM-H, Stanford University, Stanford, CA 94305, United States

### Author Contributions

†X.L. and D.S.R contributed equally to this work.

## FUNDING SOURCES

This research was supported by NIH R01 GM061238 and its successor R35 GM151985 (S.G.). D.S.R. is the Connie and Bob Lurie Fellow of the Damon Runyon Cancer Research Foundation (DRG-2526-24). D.S.R. would also like to acknowledge support from the American Heart Association Predoctoral Fellowship Grant No. 832615/David S. Roberts/2021. Y.G. would like to acknowledge support by S10 OD018475. Y.G and S.J. would like to acknowledge support by NIH R01 GM117058.

## NOTES

The authors declare no competing financial interest.

## ACKNOWLEDGMENT

We are grateful to Dr. Ilia Guzei and Dr. Zhen Yu for the consultation on X-ray crystallography.

## Notes

### Competing Interest Statement

The authors have declared no competing interest.

