## Supporting Information for "Rational design of a foldon-derived heterotrimer guided by quantitative native mass spectrometry"

### 1. Materials and Instrumentation

#### a. Materials and reagents

Amino acids and coupling reagents were obtained from Chem-Impex International, Inc. Solid-phase peptide synthesis (SPPS) resins were obtained from CEM. Sigma-Aldrich ACS-grade DMF was used as a washing solvent during SPPS, and Sigma-Aldrich biotech-grade DMF was used during amino acid coupling. Aqueous solutions were made in nanopure deionized water from Milli-Q water (MilliporeSigma). Ammonium acetate (AA, LC-MS grade) was purchased from Sigma-Aldrich (MilliporeSigma).

#### b. Instrumentation

SPPS was performed using a CEM Liberty Blue automated microwave peptide synthesizer. Preparative HPLC was performed using a Waters HPLC system (SCL-10VP system controller, LC-6AD pumps, SIL-10ADVP autosampler, SPD-10VP UV-vis detector, FRC-10A fraction collector) equipped with a Waters XSelect CSH Prep C18 column (5 µm particle size, 19 mm x 250 mm), operating at 15 mL/min. Peptide purity and concentration measurements were performed on a Waters Acquity H-Class UPLC equipped with an Acquity UPLC BEH C18 column (130 Å pore size, 1.7 µm particle size, 2.1 mm × 100 mm) operating at 0.35 mL/min. Peptide identity was determined via MALDI-TOF-MS analysis on a Bruker microflex LRF. Circular dichroism spectra were acquired on a Jasco Model J-1500 CD spectrometer. MS samples were analyzed by nanoelectrospray ionization via direct infusion using a TriVersa Nanomate system (Advion BioSciences, Ithaca, NY, USA) coupled to a solariX XR 12-T Fourier Transform Ion Cyclotron Resonance mass spectrometer (FTICR-MS, Bruker Daltonics, Billerica, MA, USA). Crystal diffraction data was collected at APS beamline 23ID-B at 0.6119Å on a Dectrics Eiger 16-M detector.

#### c. Native MS sample preparation

Native peptide samples were prepared by buffer exchanging into 150 mM ammonium acetate solution using a Zeba spin desalting column, 40K MWCO, 0.5 mL purchased from ThermoFisher Scientific. The protein sample was then diluted to 100 μM prior to native top-down MS analysis. Peptide renaturing was performed by heating peptide solutions at 95 °C for 30 min using an Eppendorf Thermomixer R Mixer (Eppendorf North America, Enfield, CT, USA), followed by cooling to 4 °C over the course of 45 min.

**d. Native top-down MS experiments**

For the nanoelectrospray ionization source using a TriVersa Nanomate, the desolvating gas pressure was set at 0.45 PSI and the voltage was set to 1.2–1.6 kV versus the inlet of the mass spectrometer. The source dry gas flow rate was set to 4 L/min at 124 °C. For the source optics, the capillary exit, deflector plate, funnel 1, skimmer voltage, funnel RF amplitude, octopole frequency, octopole RF amplitude, collision cell RF frequency, and collision cell RF amplitude were optimized at 190 V, 200 V, 100 V, 10 V, 300 Vpp, 2 MHz, 490 Vpp, 2 MHz, and 2000 Vpp, respectively. Mass spectra were acquired with an acquisition size of 4M-words of data (with a resolution of 530000 at 400 *m*/*z*) in the mass range between 200 and 4000 *m*/*z*, and a minimum of 128 scans were accumulated for each sample. Ions were accumulated in the collision cell for 10 ms, and a time-of-flight of 1.5 ms was used prior to their transfer to the ICR cell. For collisionally activated dissociation (CAD) tandem MS (MS/MS) experiments, the collision energy was varied from 10 to 30 V, ion accumulation was optimized to 400 ms, and acquisition size was 4M-words of data. Tandem mass spectra were output from the DataAnalysis 4.3 (Bruker Daltonics) software and analyzed using MASH Explorer.[1]

**e. MS data Analysis**

All data were processed and analyzed using Compass DataAnalysis 4.3 and MASH Explorer.[1] Maximum Entropy algorithm (Bruker Daltonics) was used to deconvolute all mass spectra with the instrument peak width set to 0.05 for the 12T FTICR. The sophisticated numerical annotation procedure (SNAP) peak-picking algorithm (quality factor 0.4; signal-to-noise ratio (S/N) 3.0; intensity threshold 500) was applied to determine the monoisotopic mass of all detected ions. The relative abundance for each peptide oligomeric state was determined using DataAnalysis where specific oligomeric abundances were calculated as their corresponding percentages among all of the detected oligomeric states from the native MS spectra. MS/MS data were output from the DataAnalysis software and analyzed using MASH Explorer for sequence mapping. All of the program-processed data were manually validated. All fragment ions were manually validated using MASH Explorer. Peak extraction was performed using a signal-to-noise ratio of 3 and a minimum fit of 60%, and all peaks were subjected to manual validation. All identifications were made with satisfactory numbers of assigned fragment (> 10), and a 25 ppm mass tolerance was used to match the experimental fragment ions to the calculated fragment ions based on the amino acid sequence.

**f. Single crystal formation**

All the crystals were prepared in greased 48 well VDX plate with 12 mm circle cover slides through hanging drop method. Each crystal setup contained 100 μL of buffer in the reservoir, and 1 μL peptide sample mixed with 1 μL buffer on the cover slides. The 2:1 G1K-Q11E-V14A+G1K-Q11E-V14L peptide mixture was crystalized at 2 mg/mL in 2 mM phosphate buffer, 40% w/v ethylene glycol, pH 7.5.

**g. X-ray crystallography data analysis**

The data was integrated with XDS to 0.97Å and scaled with XSCALE.[2] The structure was solved using phenix.phaser using a model derived from PDB: 1RFO.[3–5] Initially, a homotrimer model was built and refined using arp/warp and phenix.refine, interleaved with manual map fitting in Coot.[6–8] When no additional improvements were possible using a homotrimeric model, the model was adjusted to reflect the sequences of the peptides included in the experiment. The electron density maps were consistent with a heterotrimer that was rotationally disordered around the local threefold axis. It was clear that the alternative sequences at the core did not localize a specific sequence in the electron density for the three polypeptide chains. The electron density showed evidence for alternative backbone locations along the electron density for the three peptides. The most successful model used three alternative conformations for each chain, with local sequences at position 14 chain A(L,A,A), chain B(A,L,A), chain C(A,A,L). At this resolution, it could also be appreciated that the alternative chain locations also perturbed the bound solvent structure, so each alternative chain location was associated with a unique set of bound solvent. Although phenix.refine handled this model correctly, it was necessary to separate the three alternative locations and sequences for rebuilding in coot and repack the heterotrimer for additional refinement in phenix. The final model was validated using MolProbity and required files assembled with pdb_extract and deposited in the Protein Data Bank (PDB ID=8UDN).[9,10]

**h. Circular dichroism experiments and data analysis**

Circular dichroism spectra were acquired on a Jasco Model J-1500 CD spectrometer at 25.0 °C. Wavelength scans were collected from 260 to 185 nm with a 1 nm bandwidth, 0.1 nm wavelength step, and an averaging time of 4 sec per step. Concentration of each peptide was 50 μM measured by UV-Vis absorbance at 280 nm. Melting curves were acquired at 228 nm with 1.5 min equilibration at each temperature and an averaging time of 5 s. Thermal denaturation was not reversible. CD Molar Ellipticity (deg·cm$^2$·dmol$^{-1}$) was converted according to the equation[11,12]:

$$[\Theta] = \frac{\text{Ellipticity (mdeg)} \times 10^6}{\text{Pathlength (mm)} \times [\text{Protein}](\mu\text{M}) \times (\text{n})} \qquad \textit{Equation } 1$$

Where n is the number of amide bonds in the peptide.

**i. Instrument acknowledgements**

**<u>Instrument Name, Instrument Type, Grant Award Year</u>**

Crystallization and structure solution were supported by the Collaborative Crystallography Core in the Department of Biochemistry, UW-Madison.

Bruker microflex LRF, MALDI-TOF-MS, Generous gift from the Bender Fund

TriVersa Nanomate, nanoelectrospray instrument, NIH S10 OD018475

solariX XR 12-T, FTICR-MS, S10 OD018475

**2. Peptide Synthesis and Purification**

Rink Amide ProTide resin (LL) (0.19 mmol/g) was added to a CEM Liberty Blue automated microwave peptide synthesizer. Resin was swelled in DCM for 10 minutes before beginning synthesis. Fmoc amino acid (0.2 M), DIC (0.5 M), and Oxyma (1 M) were prepared and loaded to the synthesizer. 20 %v/v piperidine in ACS DMF with 0.1 M Oxyma was used as Fmoc deprotecting solvent. The coupling program was set to hold at 70°C for 10 minutes and the coupling process was doubled for Arg and β-branched amino acids. To avoid aspartimide formation with "Asp-Gly" in the sequence, backbone protected Fmoc-Asp(OtBu)-(Dmb)Gly-OH purchased from CEM was used and was coupling program was set to hold at 70°C for 20 minutes. The Fmoc deprotection program was set to hold at 90°C for 1 minute. After synthesis, the resin was dried by washing with DCM and leaving on the aspirator for 15 minutes. Cleavage was performed by adding 8 mL of 2.5 % TIPS, 2.5 % water, 95% TFA to a Torviq reaction vessel with a stir bar. The cleavage mixture was agitated in a CEM MARS microwave system at 40°C for 30 minutes, and then at room temperature for another 1.5 hours. Crude peptide solution was expunged into a 50-mL centrifuge tube, resin was washed 2X with TFA, and TFA was blown off from the combined filtrate under a stream of $N_2$. Once most of the TFA was removed, the crude peptide was precipitated by addition of 40 mL cold diethyl ether and pelleted using a centrifuge at 4000 rpm for 10 minutes. The ether supernatant was decanted, and the crude peptide solid was dried under a stream of $N_2$. Crude peptide was prepared for HPLC purification by dissolving the solid in about 2 mL of DMSO and transferring to an HPLC vial. HPLC solvent A was 0.1 % TFA in filtered and degassed Millipore $H_2O$, and solvent B was 0.1 % TFA in acetonitrile. A linear gradient of 10-50 % B was used to identify product peaks (via MALDI-TOF MS analysis of collected fractions). A flow rate of 15 mL/min was used. The column used for HPLC purification was a Waters XSelect CSH Prep C18 column (5 µm particle size, 19 mm x 250 mm).

### 3. Supplementary tables and figures

**Table S1.** Calculated mass for all the peptides.

| | Monoisotopic Mass | Average Mass |
|---|---|---|
| **Foldon** | 3079.566 | 3081.441 |
| **Q11E-V14A** | 3052.519 | 3054.372 |
| **Q11E-V14L** | 3094.566 | 3096.452 |
| **G1K-Q11E-V14G** | 3109.576 | 3111.467 |
| **G1K-V14A** | 3122.608 | 3124.509 |
| **G1K-Q11E-V14A** | 3123.592 | 3125.493 |
| **G1K** | 3150.639 | 3152.570 |
| **G1K-Q11E** | 3151.623 | 3153.547 |
| **G1K-V14L** | 3164.655 | 3166.588 |
| **G1K-Q11E-V14L** | 3165.639 | 3167.573 |
| **G1K-Q11E-V14I** | 3165.639 | 3167.573 |
| **G1K-Q11E-V14F** | 3199.623 | 3201.590 |

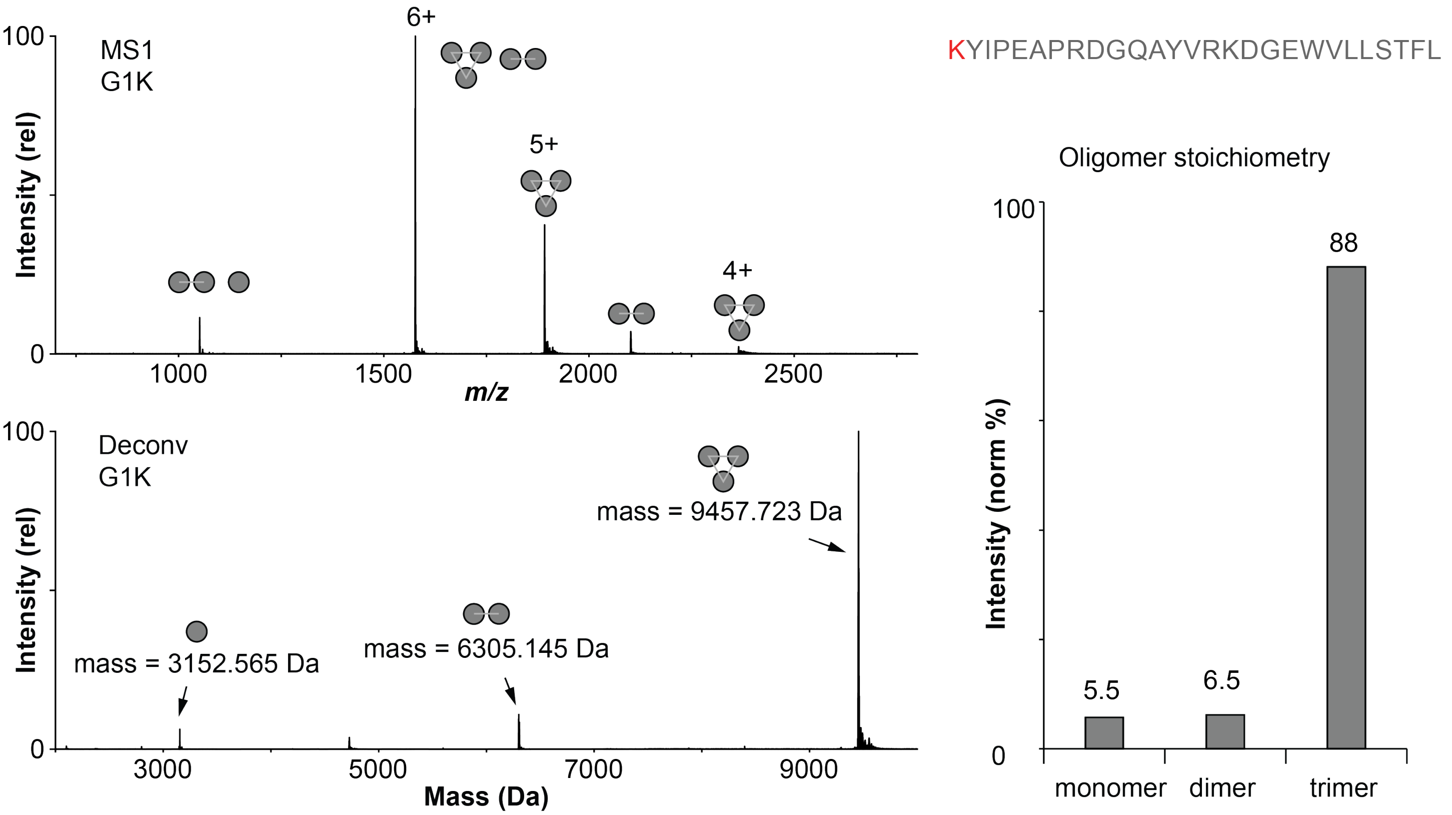


**Figure S1**. Native MS for G1K. The oligomer stoichiometry is similar to native foldon.

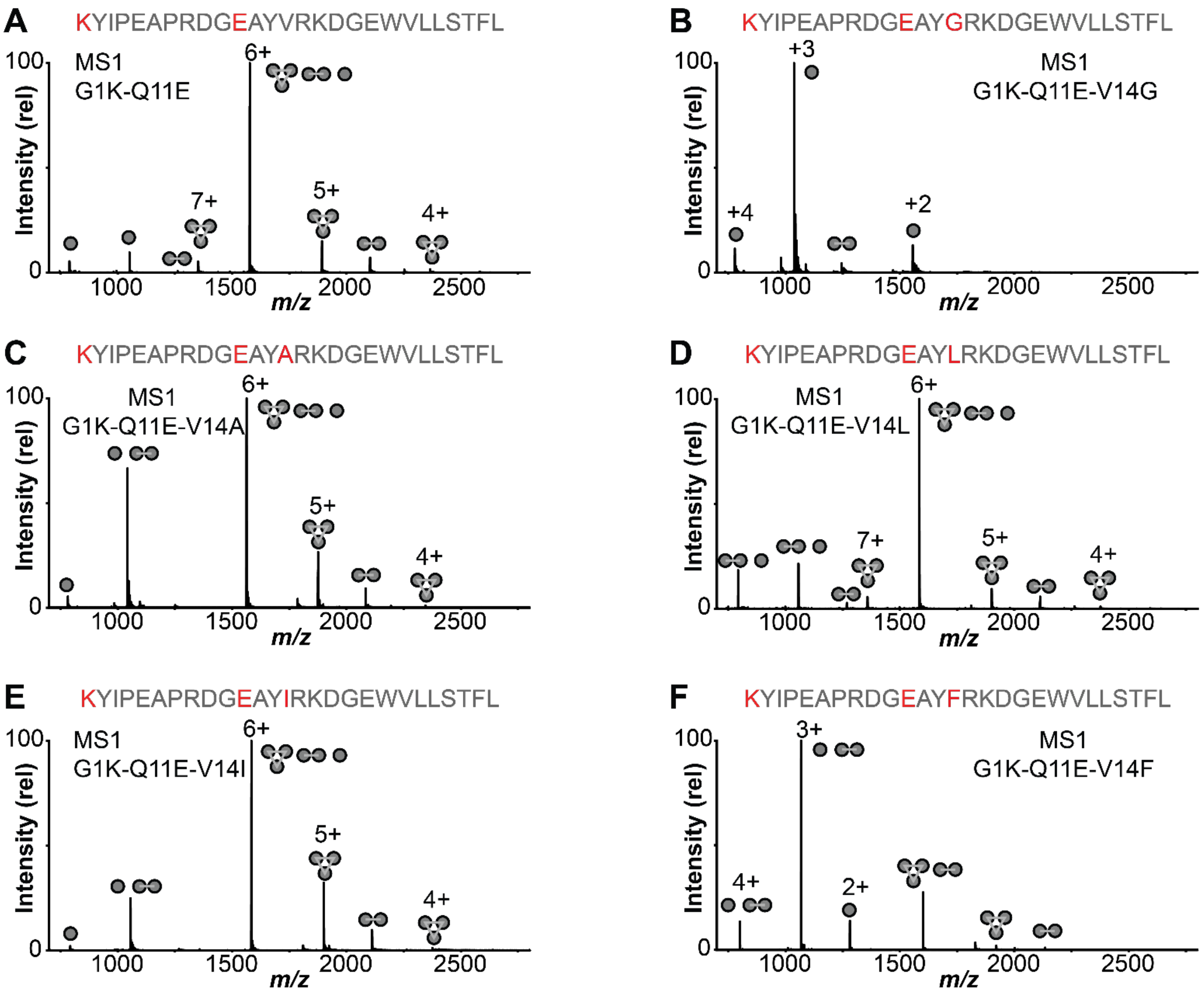


**Figure S2.** Native MS1 for all the potential heterotrimer forming peptides.

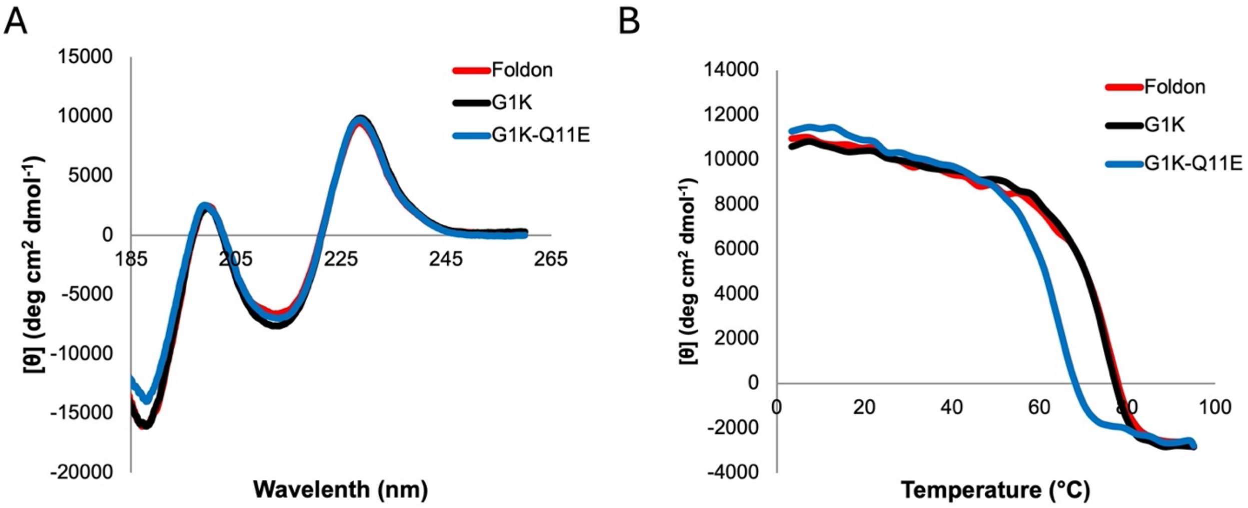


**Figure S3.** CD analysis of foldon, G1K, and G1K-Q11E. **A.** CD spectra collected in 2 mM phosphate buffer, pH=7.1 at 50 μM. **B.** Melting curve measured at 228 nm. All three peptides have similar secondary structure components, but G1K-Q11E is less thermally stable.

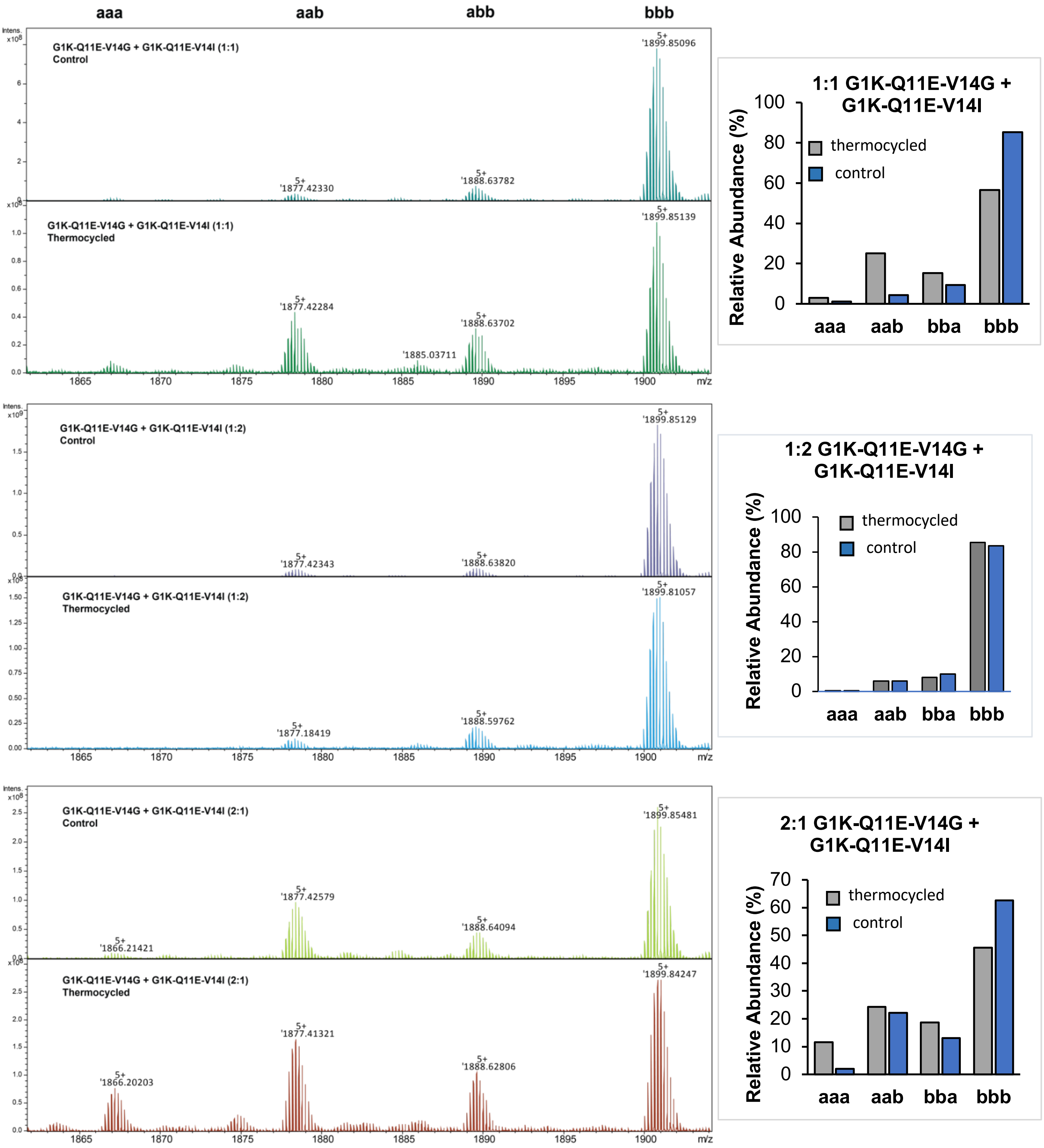


**Figure S4.** Zoomed-in native MS at charge state 5+ for trimers formed with 1:1, 1:2, and 2:1 of G1K-Q11E-V14G (a) + G1K-Q11E-V14I (b). Monoisotopic mass is labeled for each trimer signal. The mass difference between each signal, $\Delta m_{expt}$ = 56.06 Da, matches with the theoretical value, $\Delta m_{calc}$ = 56.06 Da. There is no significant bias towards a heterotrimer.

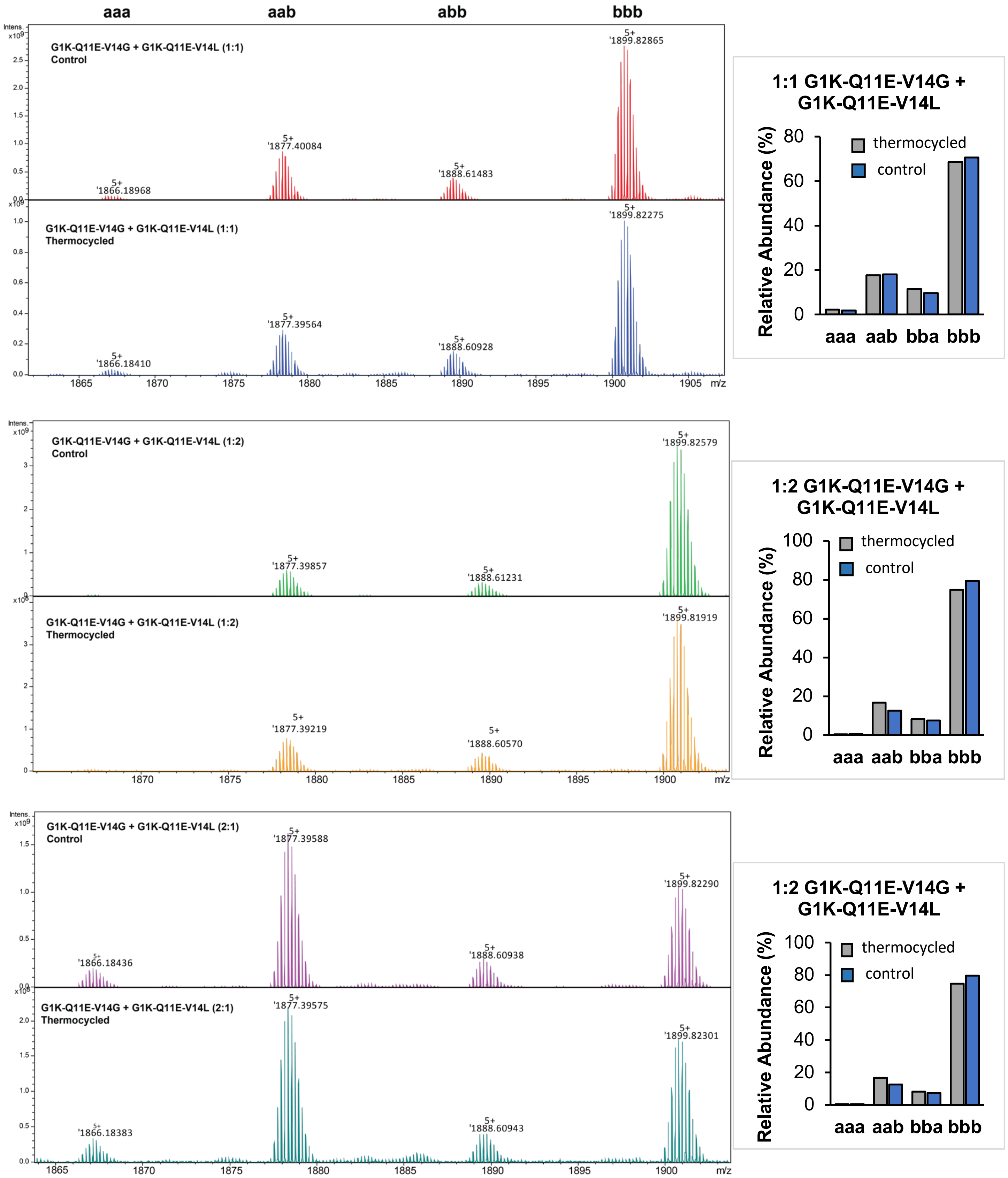


**Figure S5.** Zoomed-in native MS at charge state 5+ for trimers formed with 1:1, 1:2, and 2:1 of G1K-Q11E-V14G (a) + G1K-Q11E-V14L (b). Monoisotopic mass is labeled for each trimer signal. The mass difference between each signal, $\Delta m_{expt}$ = 56.06 Da, matches with the theoretical value, $\Delta m_{calc}$ = 56.06 Da. There is no significant bias towards a heterotrimer.

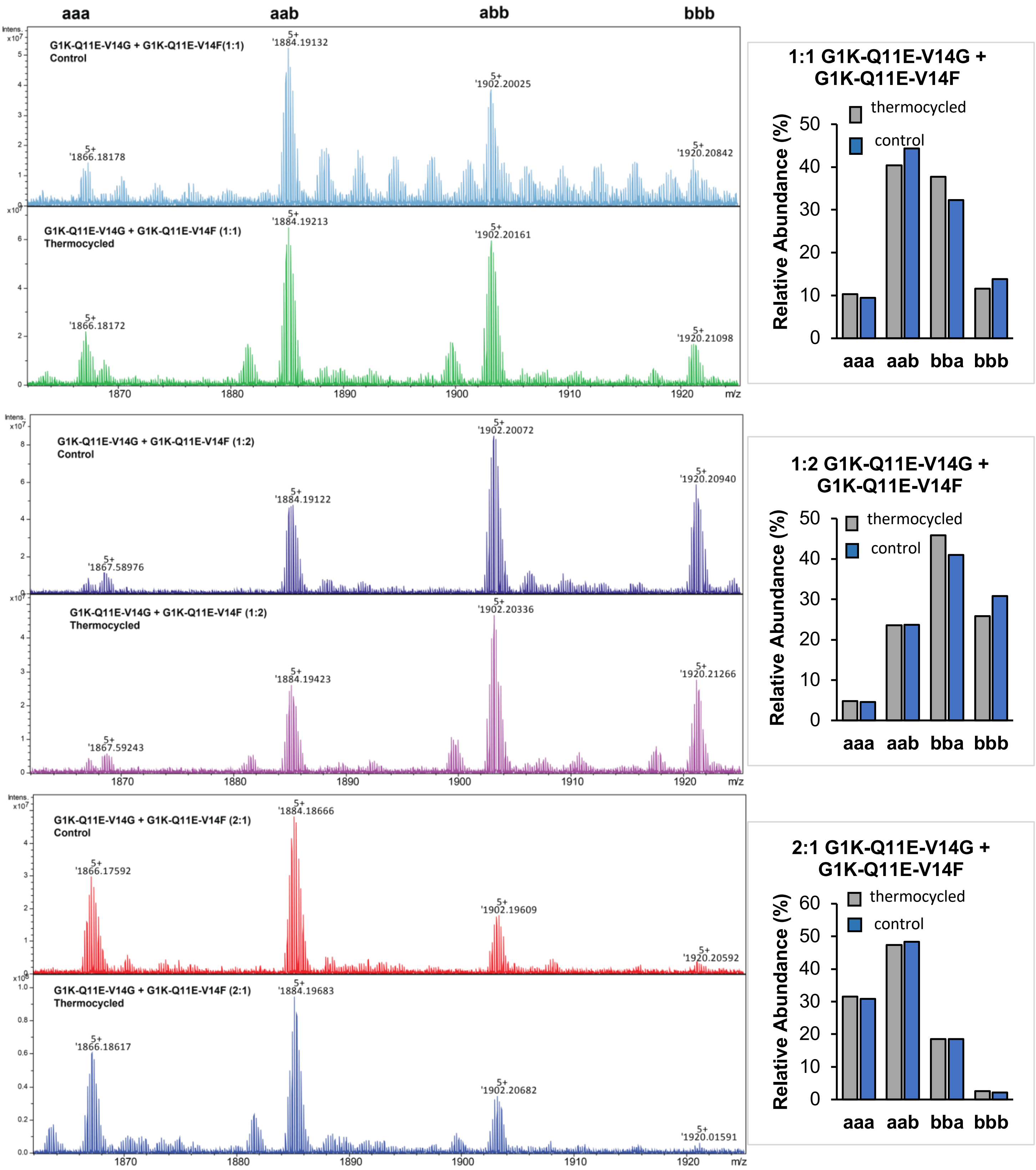


**Figure S6.** Left: Zoomed-in native MS at charge state 5+ for trimers formed with 1:1, 1:2, and 2:1 of G1K-Q11E-V14G (a) + G1K-Q11E-V14F (b). Monoisotopic mass is labeled for each trimer signal. The mass difference between each signal, $\Delta m_{expt}$ = 90.05 Da, matches with the theoretical value, $\Delta m_{calc}$ = 90.05 Da. There is no significant bias towards a heterotrimer.

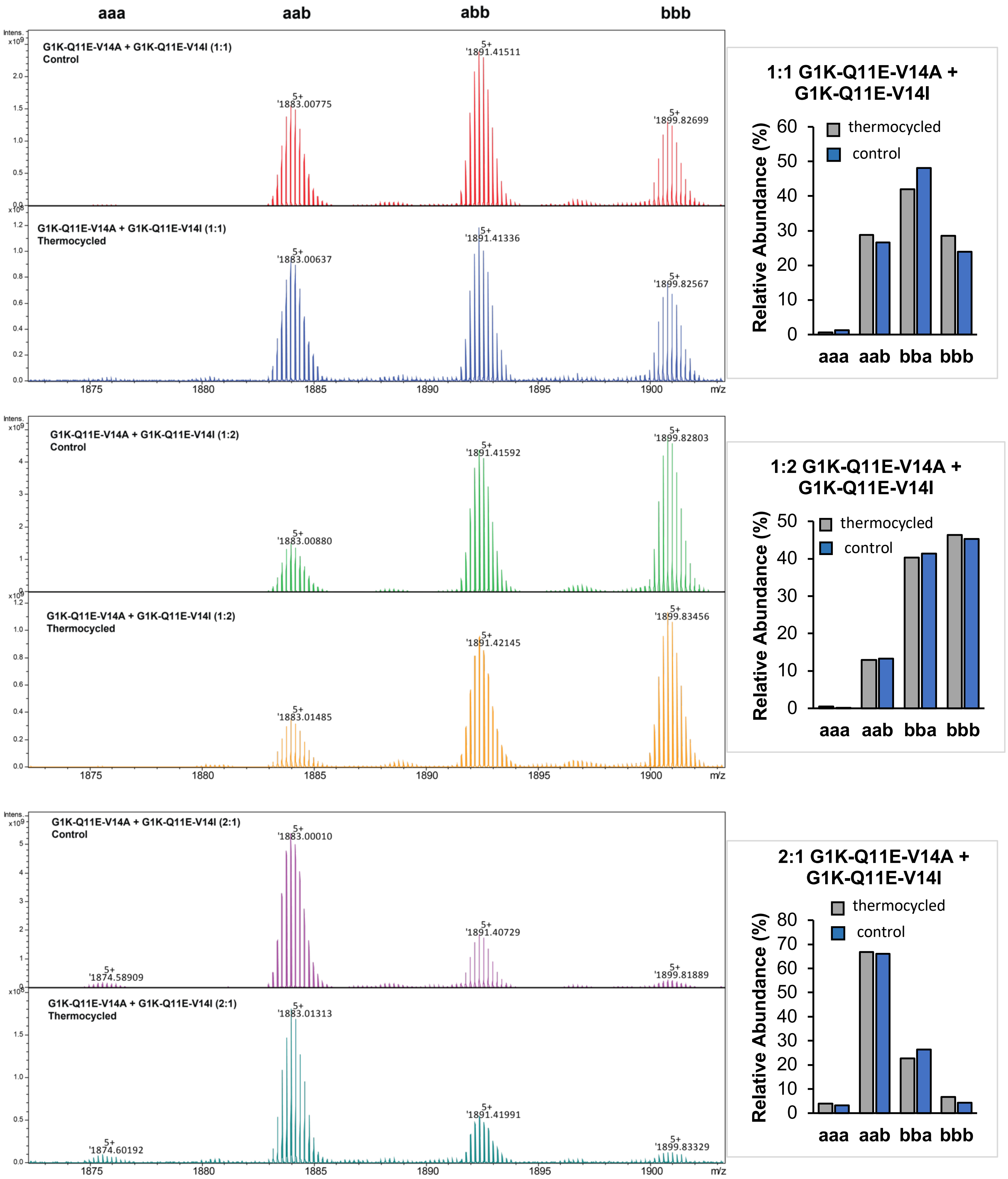


**Figure S7.** Zoomed-in native MS at charge state 5+ for trimers formed with 1:1, 1:2, and 2:1 of G1K-Q11E-V14A (a) + G1K-Q11E-V14I (b). Monoisotopic mass is labeled for each trimer signal. The mass difference between each signal, $\Delta m_{expt}$ = 42.05 Da, matches with the theoretical value, $\Delta m_{calc}$ = 42.06 Da. There is a bias towards an *aab* heterotrimer in the 2:1 mixture, but also >20% *abb* heterotrimer.

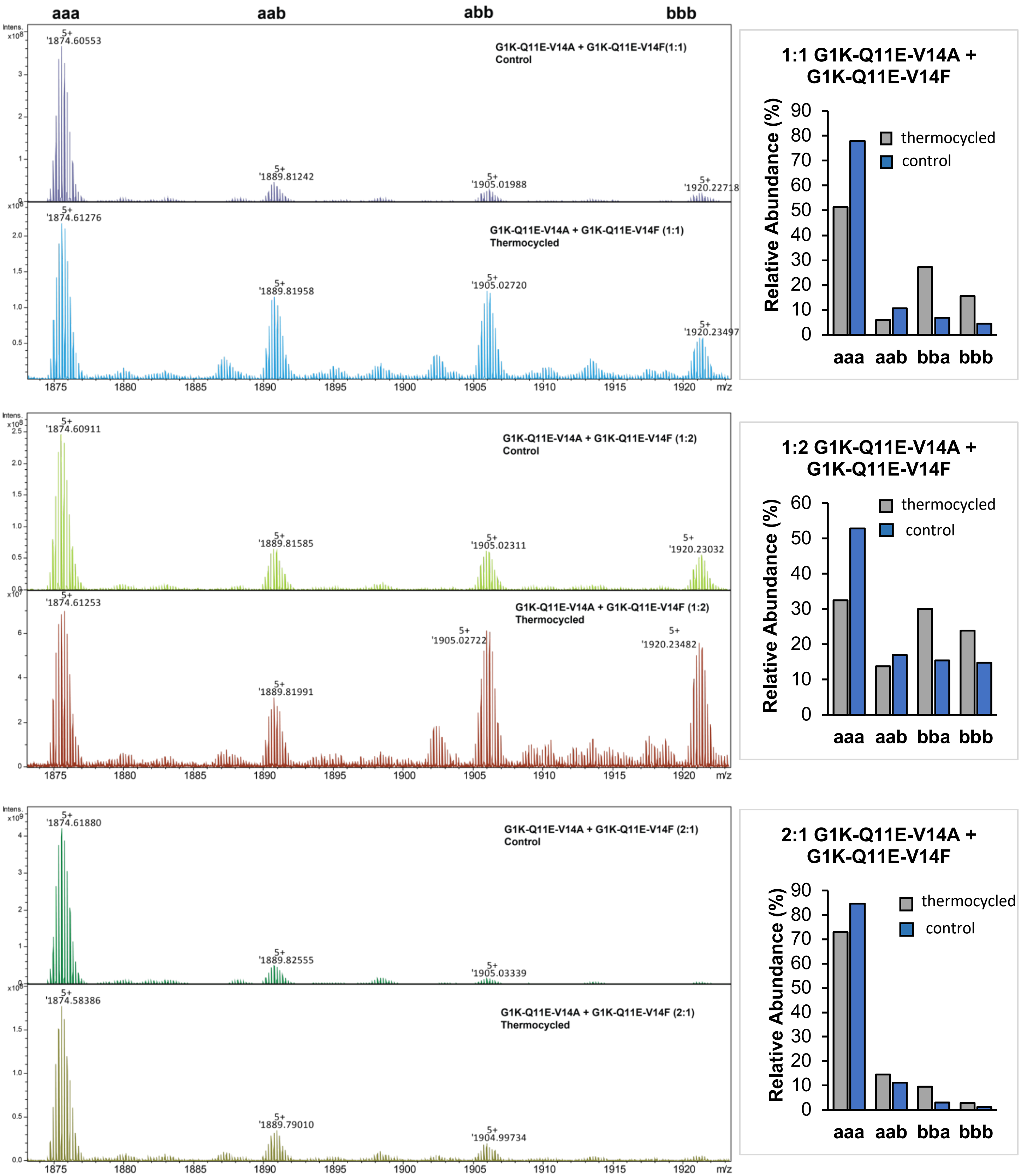


**Figure S8.** Left: Zoomed in native MS at charge state 5+ for trimers formed with 1:1, 1:2, and 2:1 of G1K-Q11E-V14A (a) + G1K-Q11E-V14F (b). Monoisotopic mass is labeled for each trimer signal. The mass difference between each signal, $\Delta m_{expt}$ = 76.03 Da, matches with the theoretical value, $\Delta m_{calc}$ = 76.03 Da. There is no significant bias towards a heterotrimer.

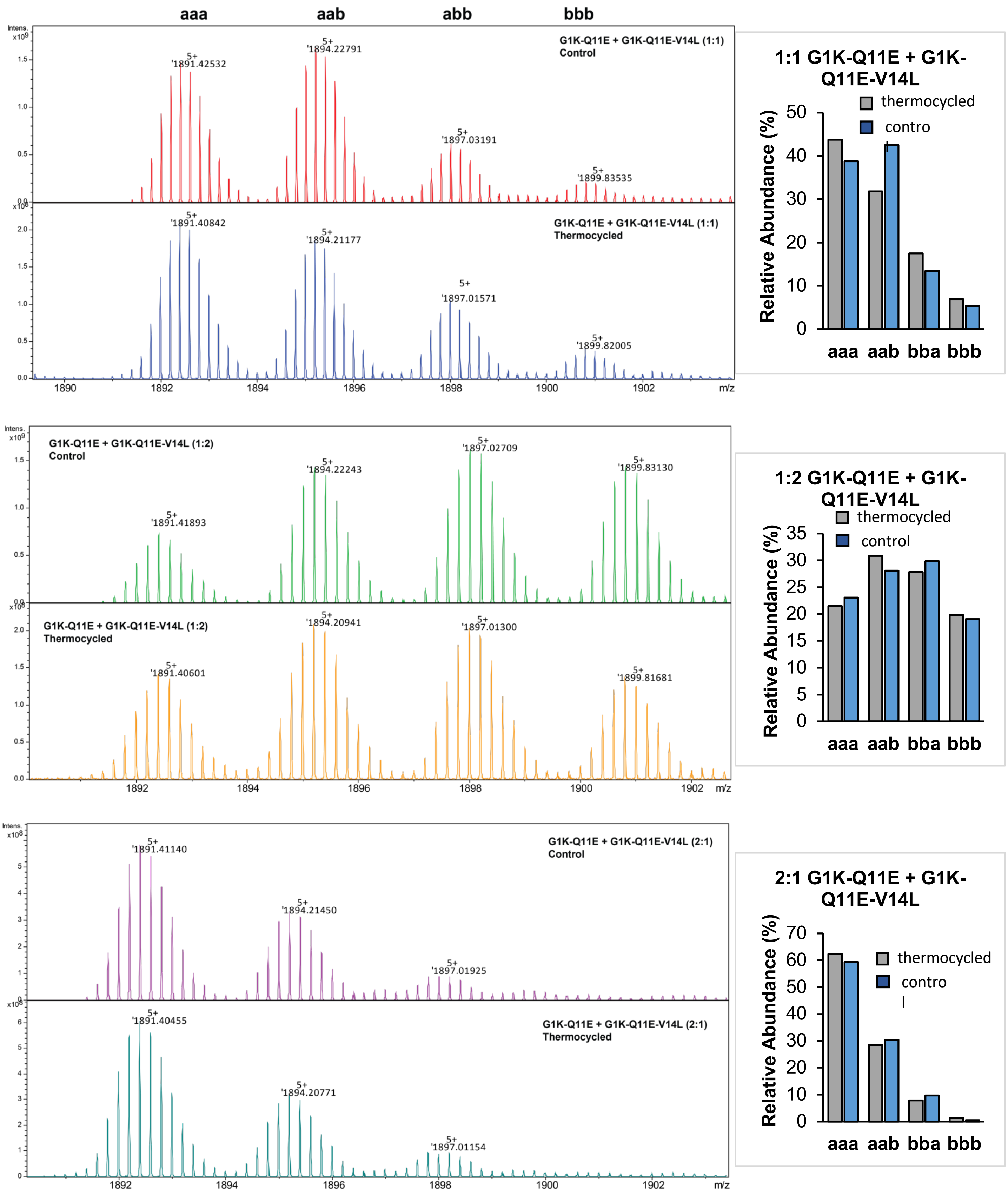


**Figure S9.** Zoomed in native MS at charge state 5+ for trimers formed with 1:1, 1:2, and 2:1 of G1K-Q11E (a) + G1K-Q11E-V14L (b). Monoisotopic mass is labeled for each trimer signal. The mass difference between each signal, $\Delta m_{expt}$ = 14.02 Da, matches with the theoretical value, $\Delta m_{calc}$ = 14.02 Da. There is no significant bias towards a heterotrimer.

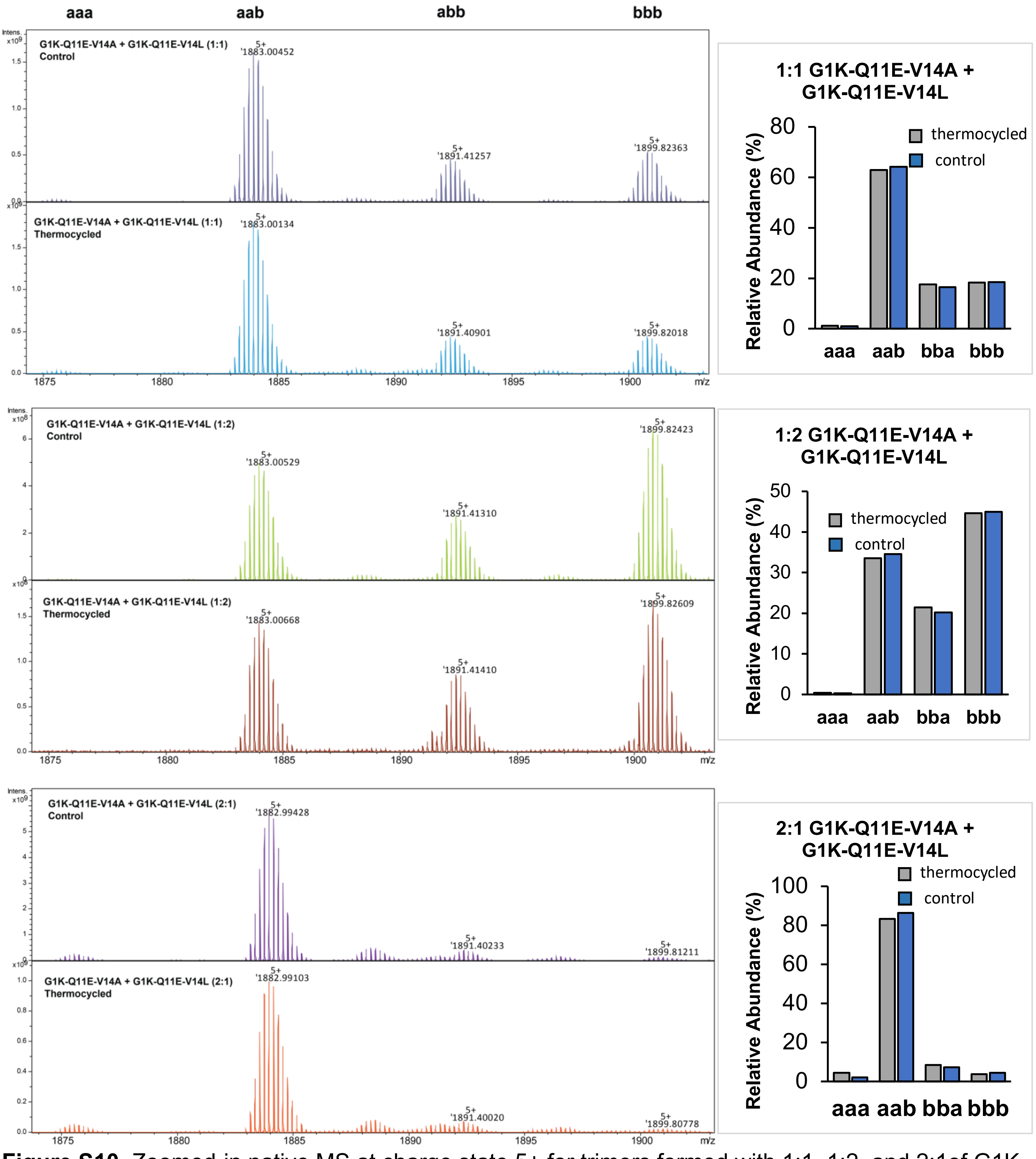


**Figure S10.** Zoomed-in native MS at charge state 5+ for trimers formed with 1:1, 1:2, and 2:1of G1K-Q11E-V14A (a) + G1K-Q11E-V14L (b). Monoisotopic mass is labeled for each trimer signal. The mass difference between each signal, $\Delta m_{expt}$ = 42.08 Da, matches with the theoretical value, $\Delta m_{calc}$ = 42.05 Da. The 2:1 stoichiometry generates a dominant *aab* heterotrimer.

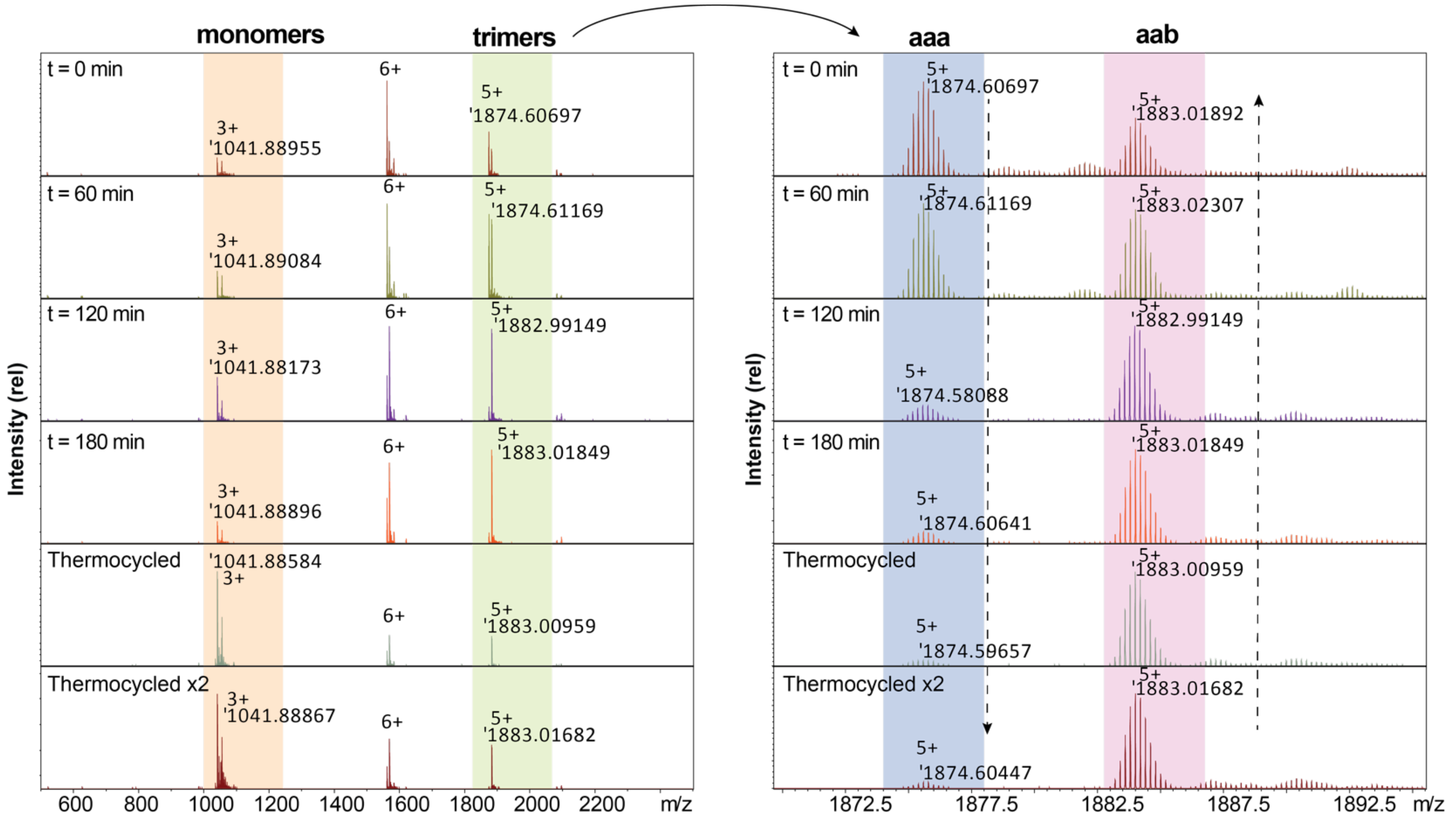


**Figure S11.** Assessment of thermocycling and mixing of 2:1 G1K-Q11E-V14A + G1K-Q11E-V14L using native MS1 and their zoomed-in spectra in the 5+ charge state trimer region. The first 4 spectra show that G1K-Q11E-V14A and G1K-Q11E-V14L were mixed as separate solutions. The mixture reached an equilibrium after 180 min at room temperature, with a dominant *aab* heterotrimer. After reaching an equilibrium, the sample was thermocycled and the native MS is shown in the "Thermocycled" spectrum, where more dissociation into monomer was observed (higher intensity on the leftmost 3+ charge state signal, which corresponds to monomeric peptides). The last spectrum "Thermocycled x2" shows two separated solution samples that were heated separately, combined, and then heated again. All the samples showed partially dissociated monomers, which increase upon heating, but the dominant trimer remains to be the *aab* heterotrimer.

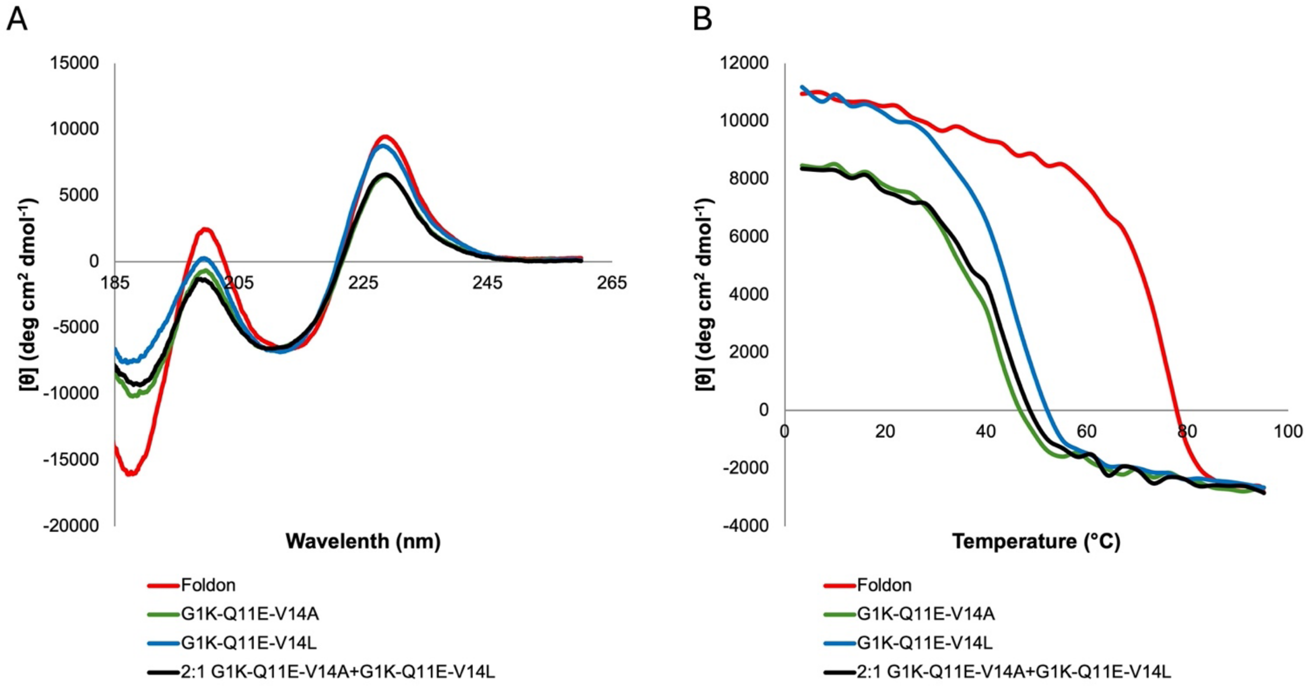


**Figure S12**. CD analysis of foldon, G1K-Q11E-V14A, G1K-Q11E-V14L, and 2:1 G1K-Q11E-V14A + G1K-Q11E-V14L. **A.** CD spectra collected in 2 mM phosphate buffer, pH=7.1 at 50 μM. **B.** Melting curve measured at 228 nm. The thermostability of the mixed peptides falls between the two isolated peptides.

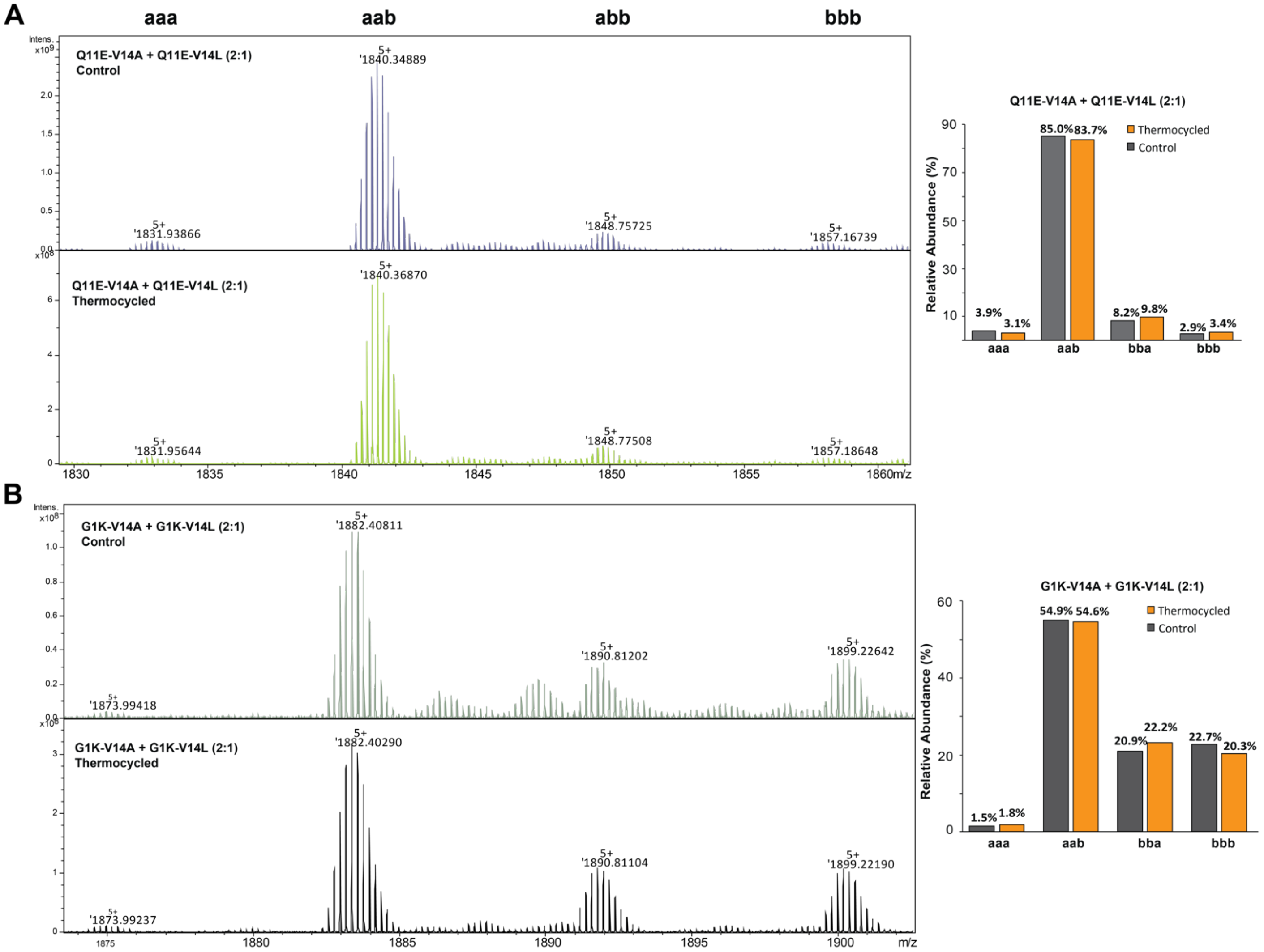


**Figure S13**. Reversed stepwise mutation to study the heterotrimerization mechanism. Zoomed-in native MS at charge state 5+ for trimers formed with **A.** 2:1 Q11E-V14A (a) + Q11E-V14L (b), and **B.** with 2:1 G1K-V14A (a) + G1K-V14L (b). Monoisotopic mass is labeled for each trimer signal. In both cases, the mass difference between each signal, $\Delta m_{expt}$ = 42.05 Da, matches with the theoretical value, $\Delta m_{calc}$ = 42.05 Da. G1K mutation did not affect the selectivity towards the *aab* heterotrimer, whereas Q11E mutation was essential for the heterotrimer selectivity.

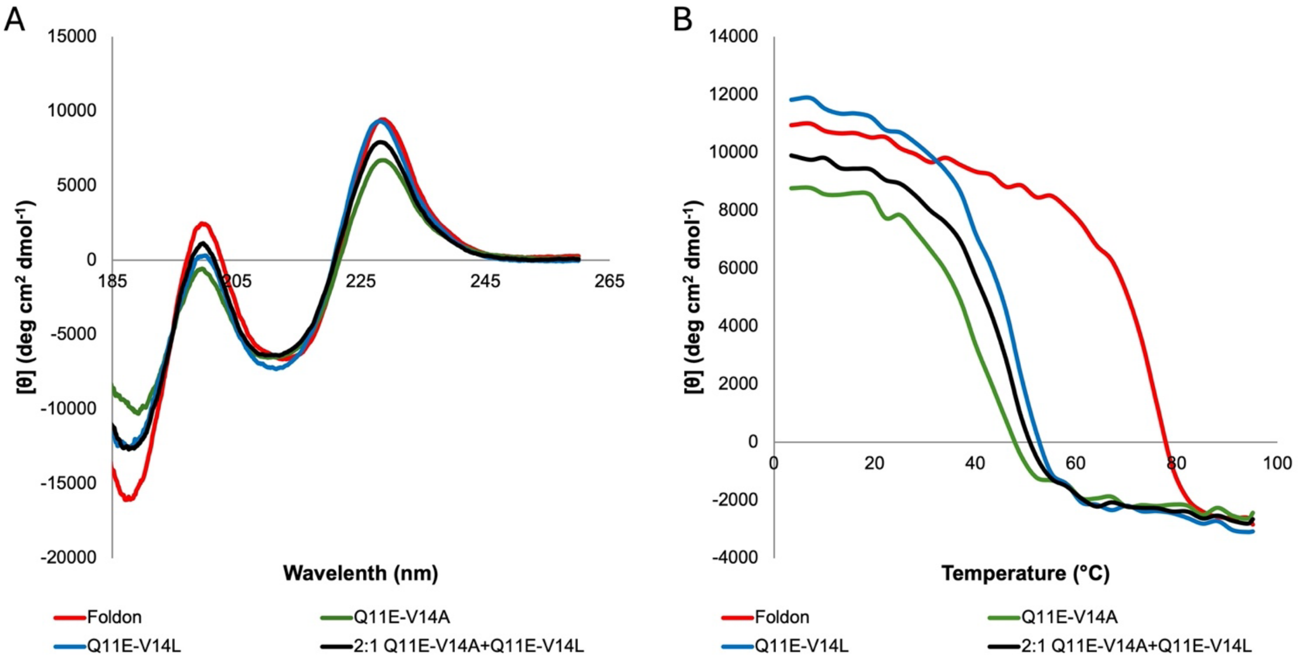


**Figure S14**. CD analysis of foldon, Q11E-V14A, Q11E-V14L, and 2:1 Q11E-V14A + Q11E-V14L. **A.** CD spectra collected in 2 mM phosphate buffer, pH=7.1 at 50 μM. **B.** Melting curve measured at 228 nm. The thermostability of the mixed peptides falls between the two isolated peptides.

**Table S2.** Melting temperature ($T_m$ ) of selected peptides and their binary mixtures.

| Peptide | $T_m$ (K) |
|---|---|
| Foldon | 348 |
| G1K | 348 |
| G1K-Q11E | 337 |
| G1K-Q11E-V14A | 314 |
| G1K-Q11E-V14L | 318 |
| 2:1 G1K-Q11E-V14A + G1K-Q11E-V14L | 316 |
| Q11E-V14A | 314 |
| Q11E-V14L | 320 |
| 2:1 Q11E-V14A + Q11E-V14L | 319 |

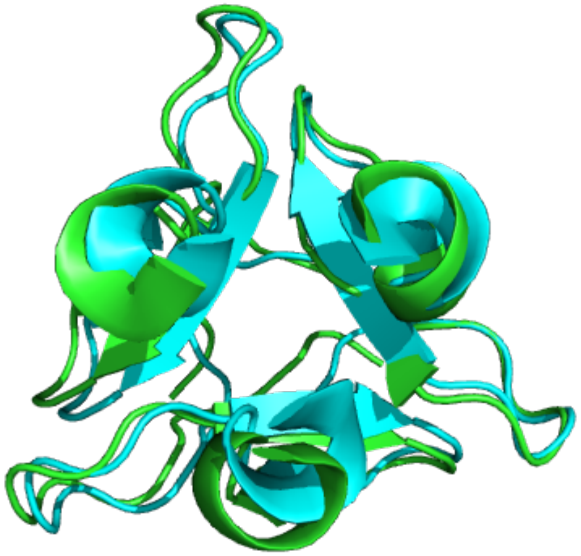

**Figure S15.** Superimposition of the crystal structure of 2:1 G1K-Q11E-V14A + G1K-Q11E-V14L (PDB: 8UDN, green), and NMR structure of native foldon (PDB: 1RFO [5], cyan).

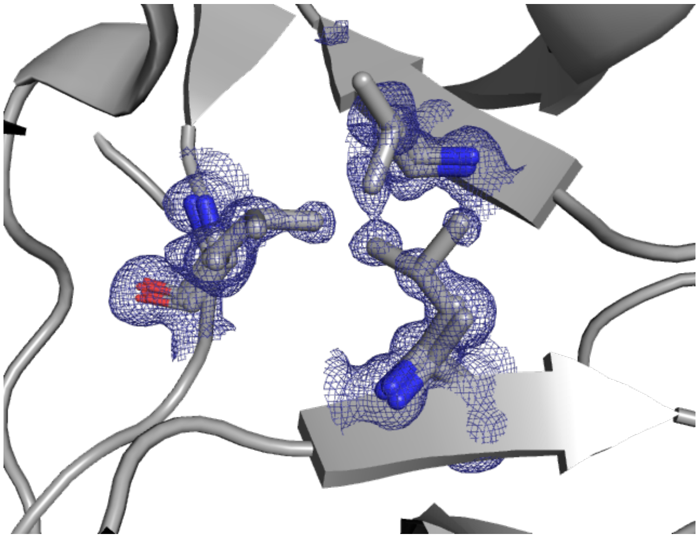

**Figure S16.** Zoomed-in electron density map at the central hydrophobic cores from the crystal structures of 2:1 G1K-Q11E-V14A + G1K-Q11E-V14L. Because of the rotationally disorder about the non-crystallographic three-fold axis. The amino acids at position 14 are depicted as three overlapping side chains with two Ala and one Leu. The electron density map of the three side chains showed a higher degree of disorder compared the rest of the structure but matches with our expectation when Leu is present at 1/3 occupancy.

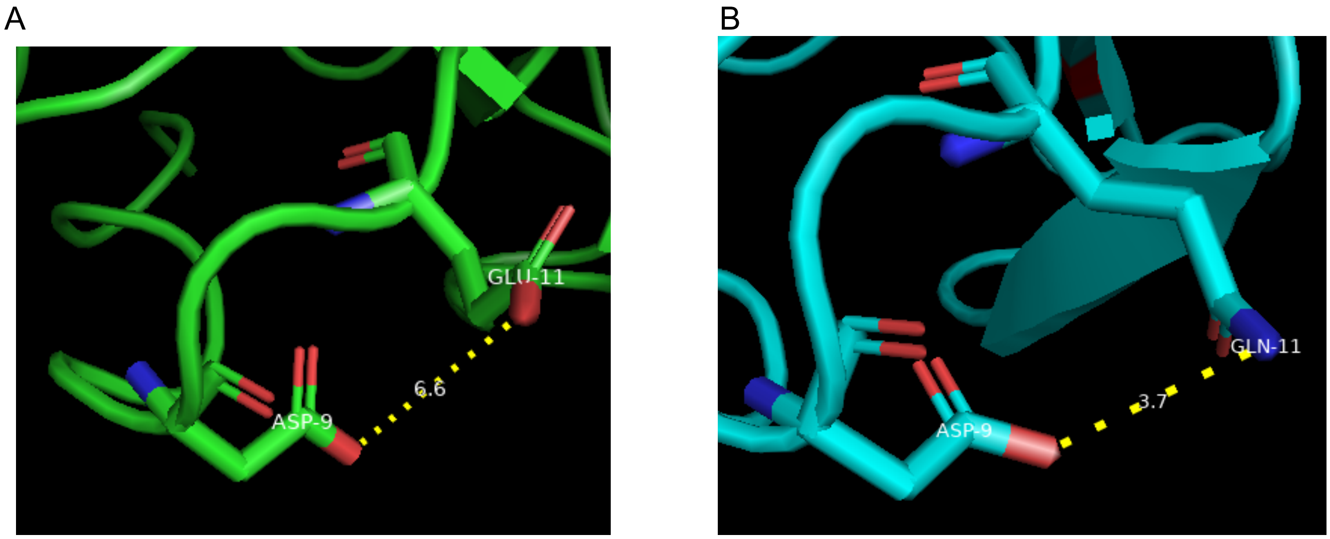


**Figure S17.** The relative positions of the amino acid side chains at position 9 and 11 in the heterotrimer is significantly longer than in the native foldon, indicating the disturbed monomeric structure from mutation Q11E. **A.** Distance between Asp 9 and Glu11 in 8UDN (6.6 Å). **B.** Distance between Asp 9 and Gln11 in 1RFO[5] (3.7 Å).

**Table S3.** X-ray diffraction data collection and refinement statistics.

| | **8UDN** |
|---|---|
| **Wavelength** | 0.6199 |
| **Resolution range** | 22.2 - 0.97 (1.005 - 0.97) |
| **Space group** | P 43 |
| **Unit cell** | 32.854 32.854 75.23 90 90 90 |
| **Total reflections** | 651233 (65258) |
| **Unique reflections** | 46606 (4623) |
| **Multiplicity** | 14.0 (14.1) |
| **Completeness (%)** | 99.04 (98.30) |
| **Mean I/sigma(I)** | 17.81 (1.37) |
| **Wilson B-factor** | 10.31 |
| **R-merge** | 0.05849 (1.196) |
| **R-meas** | 0.06067 (1.24) |
| **R-pim** | 0.016 (0.3264) |
| **CC1/2** | 1 (0.77) |
| **CC*** | 1 (0.933) |
| **Reflections used in refinement** | 46594 (4618) |
| **Reflections used for R-free** | 2006 (206) |
| **R-work** | 0.1451 (0.2606) |
| **R-free** | 0.1571 (0.2705) |
| **CC(work)** | 0.974 (0.889) |
| **CC(free)** | 0.980 (0.873) |
| **Number of non-hydrogen atoms** | 2358 |
| **macromolecules** | 2007 |
| **ligands** | 0 |
| **solvent** | 351 |
| **Protein residues** | 81 |
| **RMS(bonds)** | 0.010 |
| **RMS(angles)** | 1.10 |
| **Ramachandran favored (%)** | 94.67 |
| **Ramachandran allowed (%)** | 5.33 |
| **Ramachandran outliers (%)** | 0.00 |
| **Rotamer outliers (%)** | 0.00 |
| **Clashscore** | 6.79 |
| **Average B-factor** | 12.97 |
| **macromolecules** | 12.34 |
| **solvent** | 16.57 |
| **Number of TLS groups** | 6 |

Statistics for the highest-resolution shell are shown in parentheses.

### 4. Peptide Characterization by UPLC and MALDI-MS

UPLC settings: Waters Acquity UPLC; Waters Acquity UPLC BEH C18 column, 100 mm. 2.1 mm, 1.7 Å particle size, 130 Å pore size, general method: 10-90% B over 7 min, 0.35 mL/min. Detected channel: 280 nm.

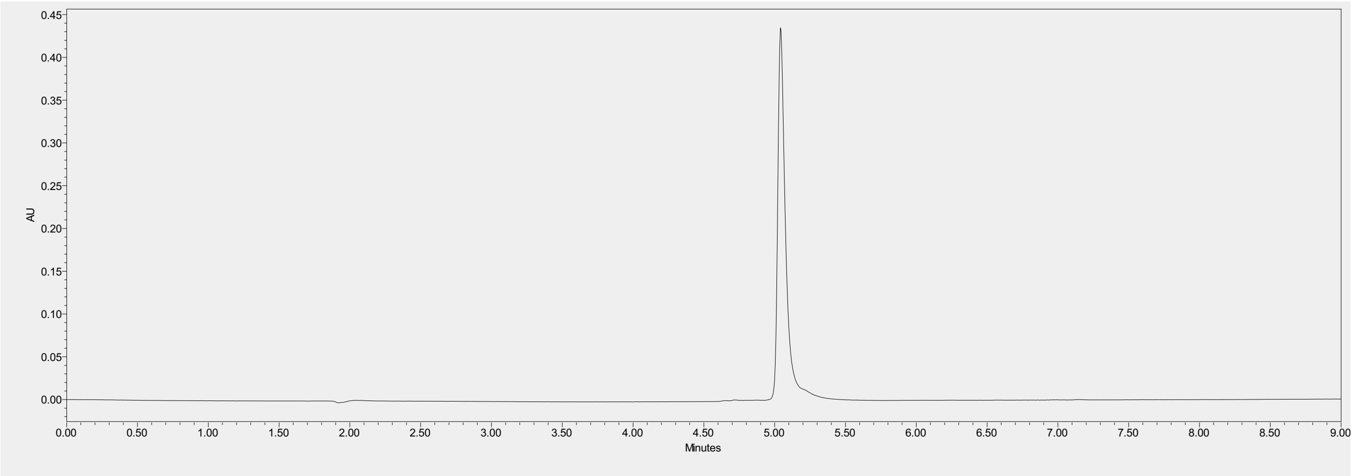


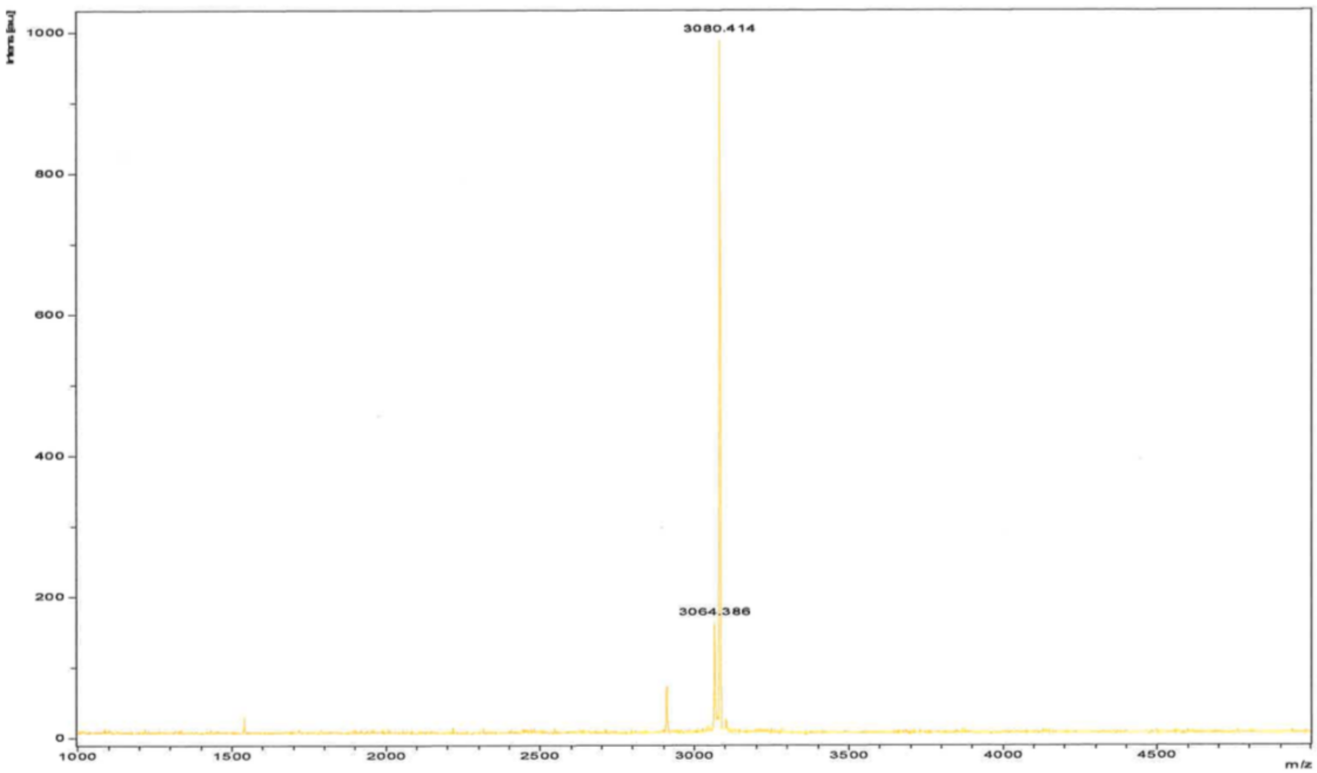


UPLC trace and MALDI-TOF MS for foldon. Peptide had purity higher than 98%. The signal at lower m/z is impurity due to aspartimide formation.

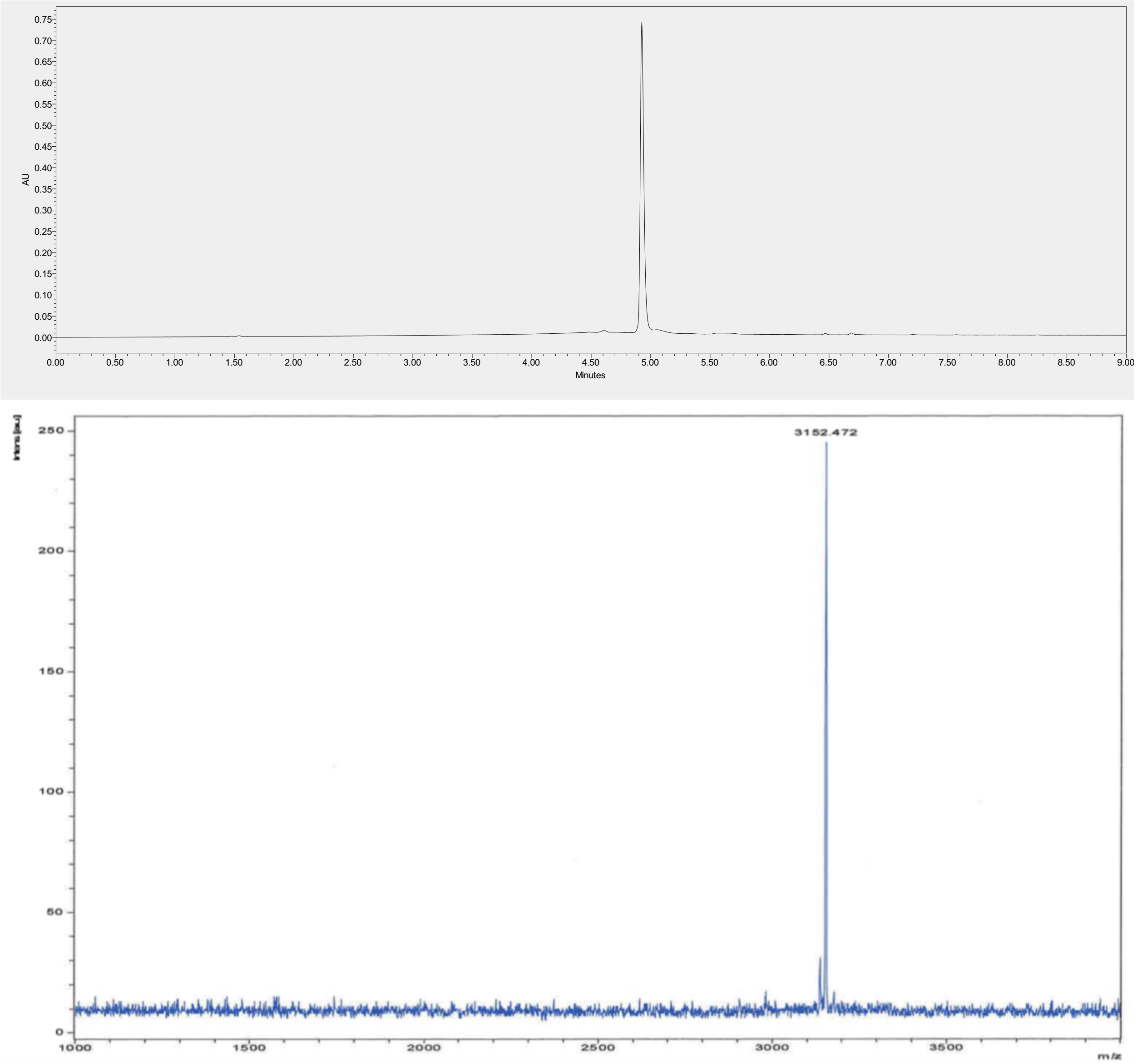


UPLC trace and MALDI-TOF MS for G1K. Peptide had purity higher than 98%.

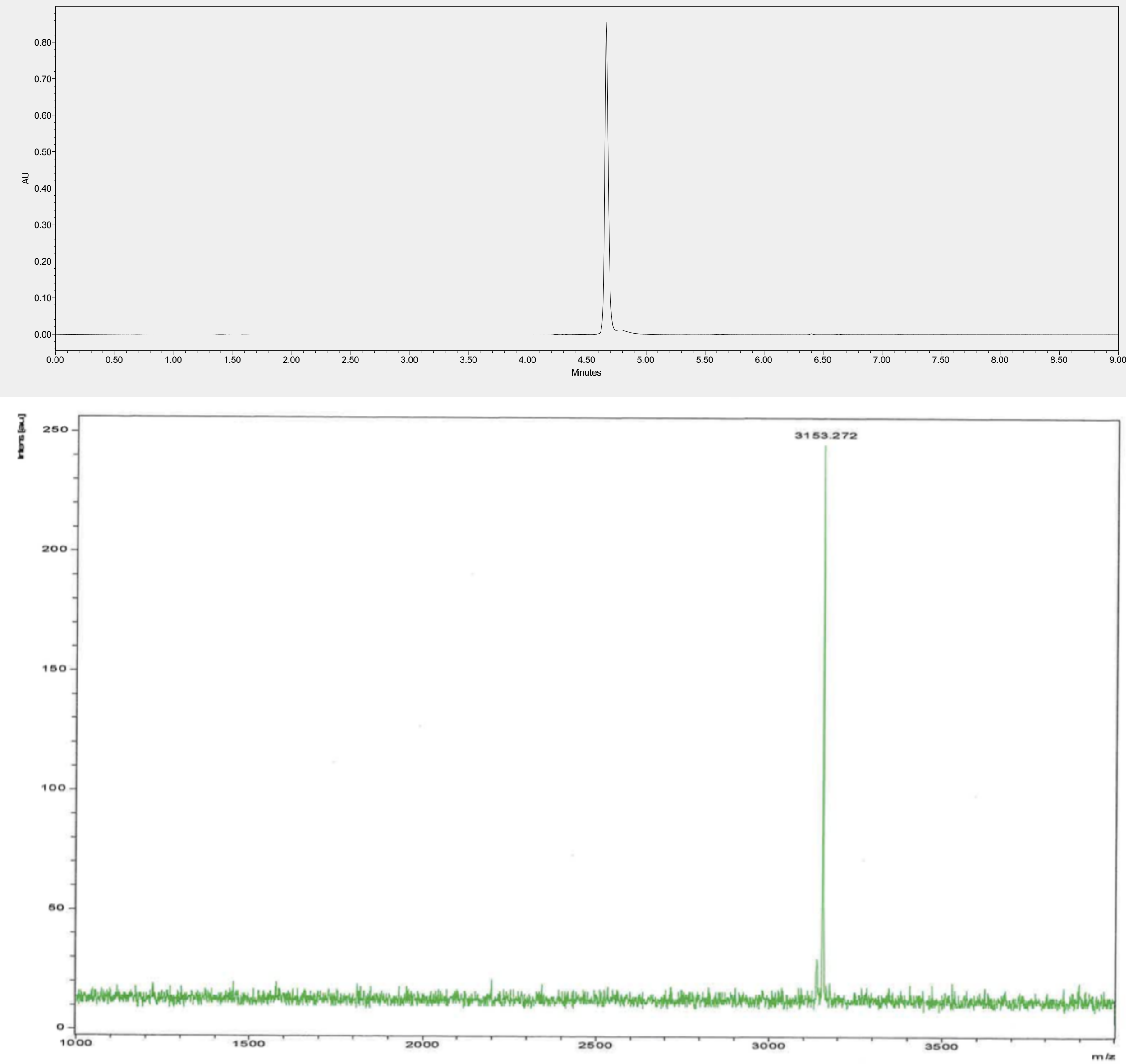


UPLC trace and MALDI-TOF MS for G1K-Q11E. Peptide had purity higher than 98%.

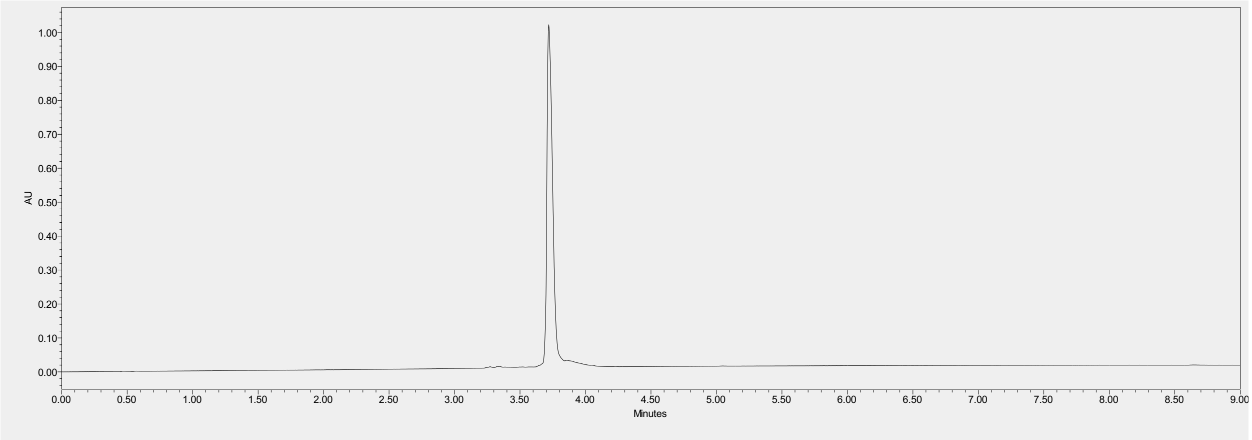


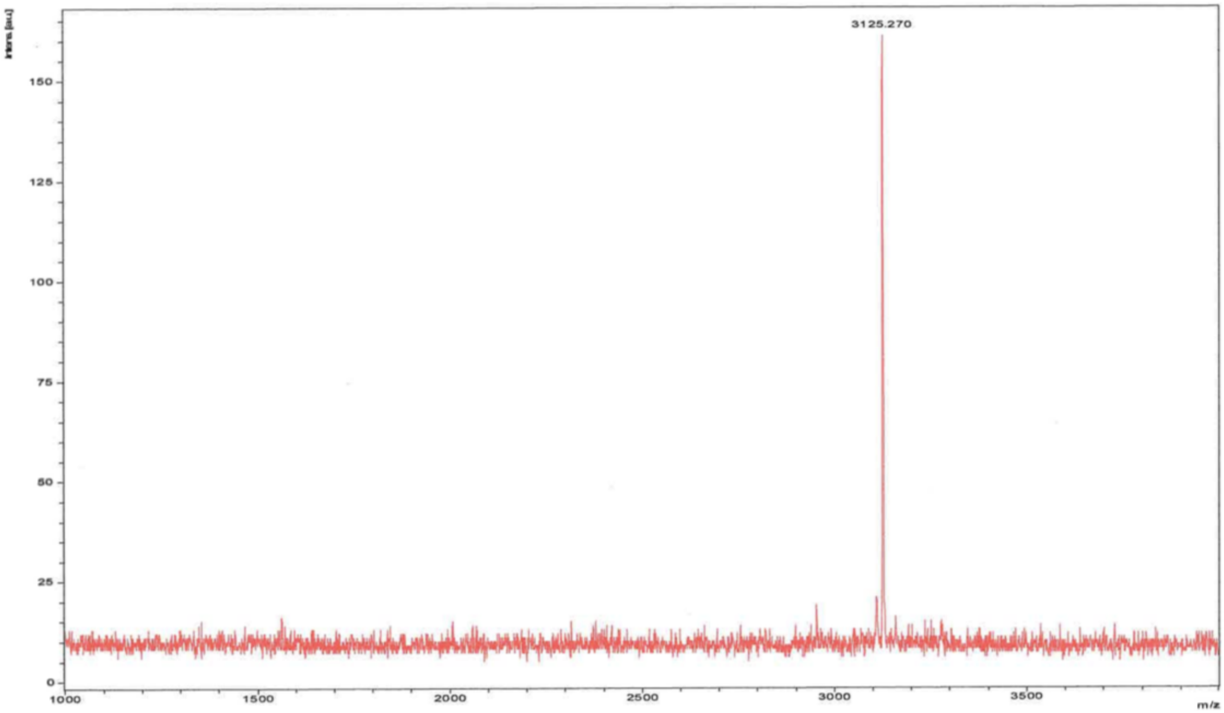


UPLC trace and MALDI-TOF MS for G1K-V14A. Peptide had purity higher than 98%.

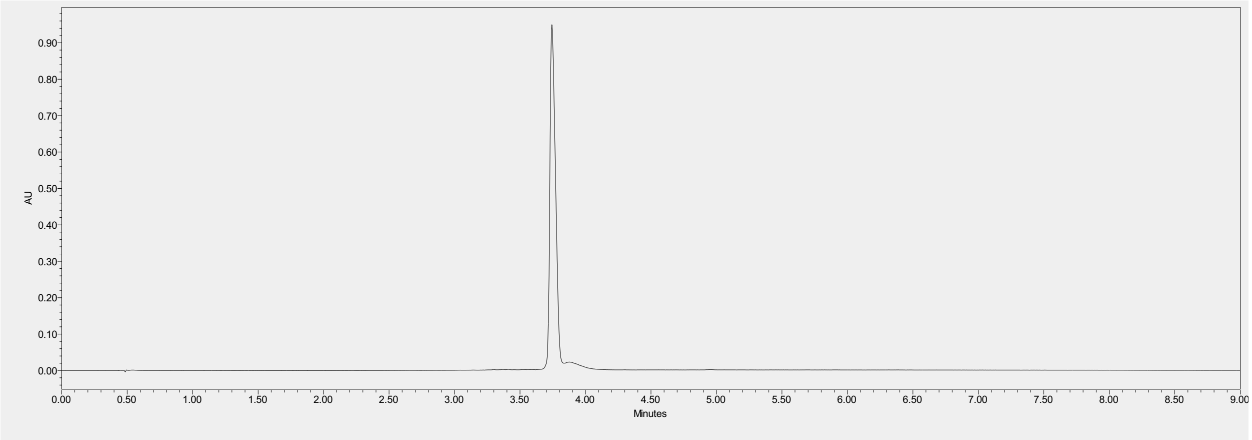


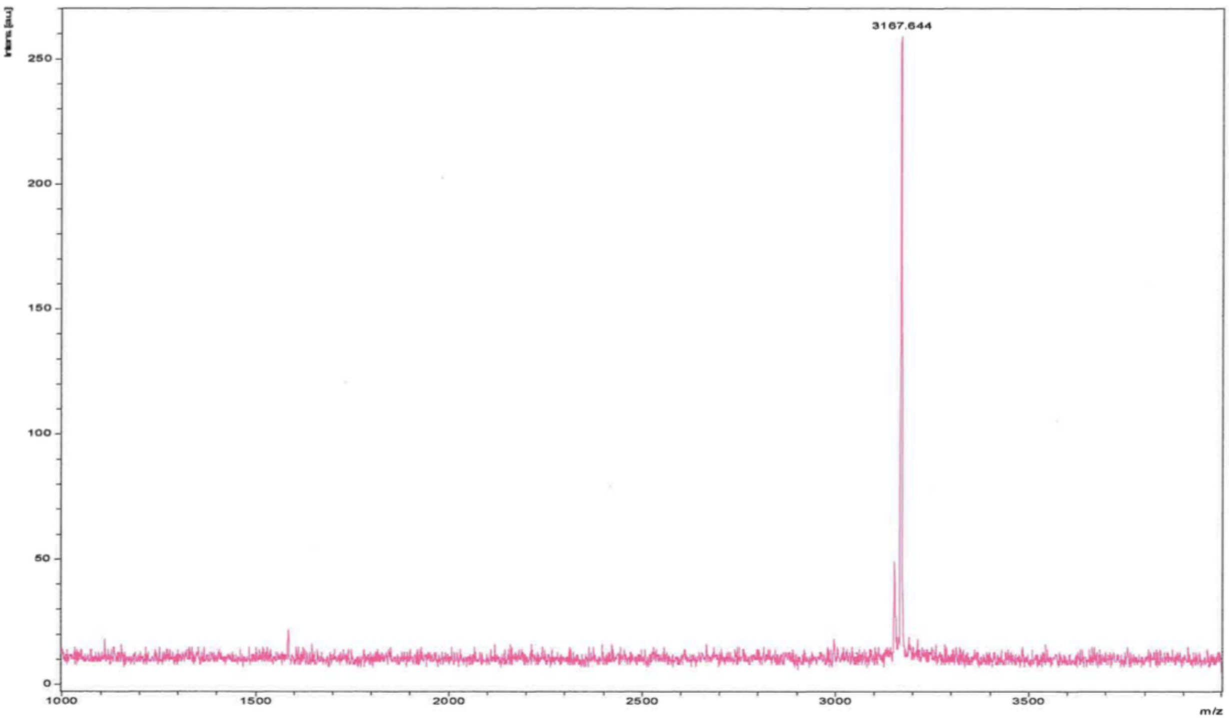


UPLC trace and MALDI-TOF MS for G1K-V14L. Peptide had purity higher than 98%.

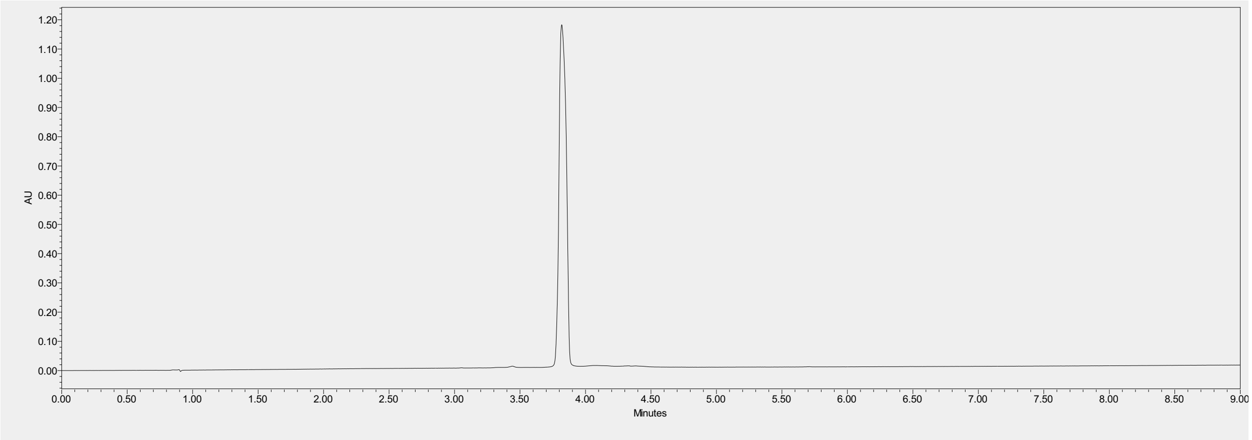


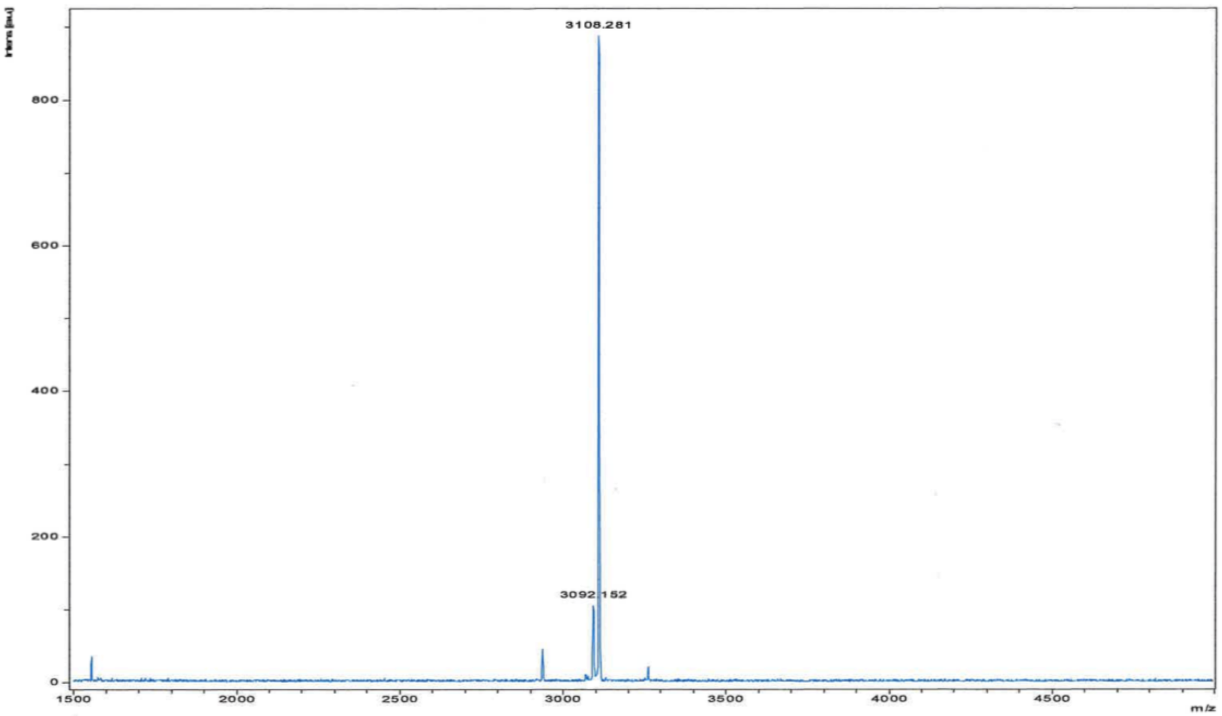


UPLC trace and MALDI-TOF MS for G1K-Q11E-V14G. Peptide had purity higher than 98%. The signal at lower m/z is impurity due to aspartimide formation.

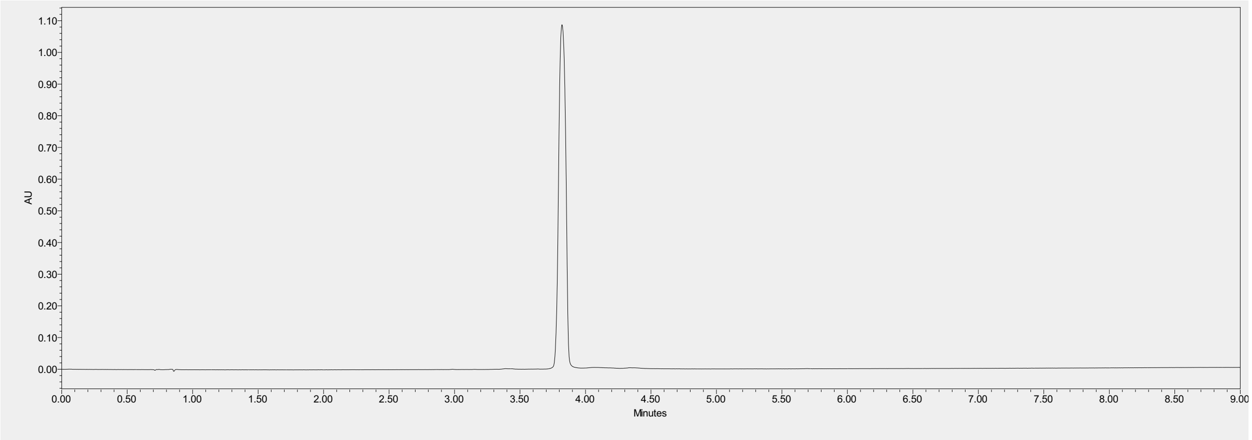


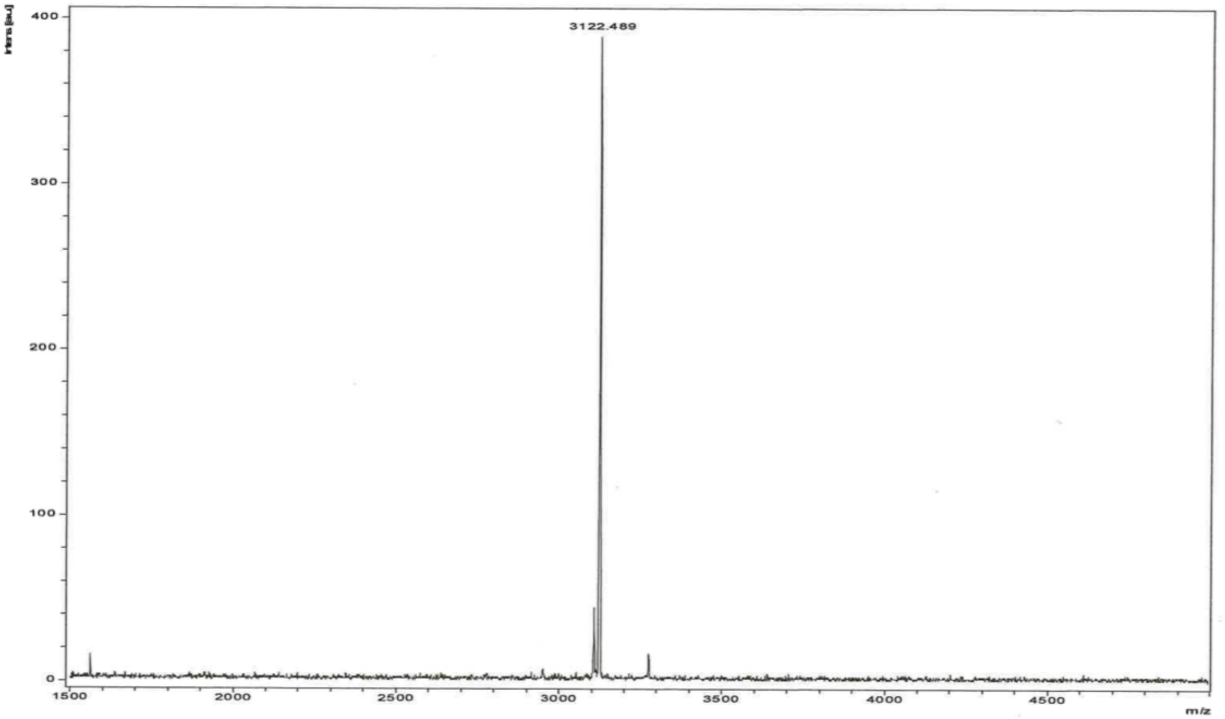


UPLC trace and MALDI-TOF MS for G1K-Q11E-V14A. Peptide had purity higher than 98%. The signal at lower m/z is impurity due to aspartimide formation.

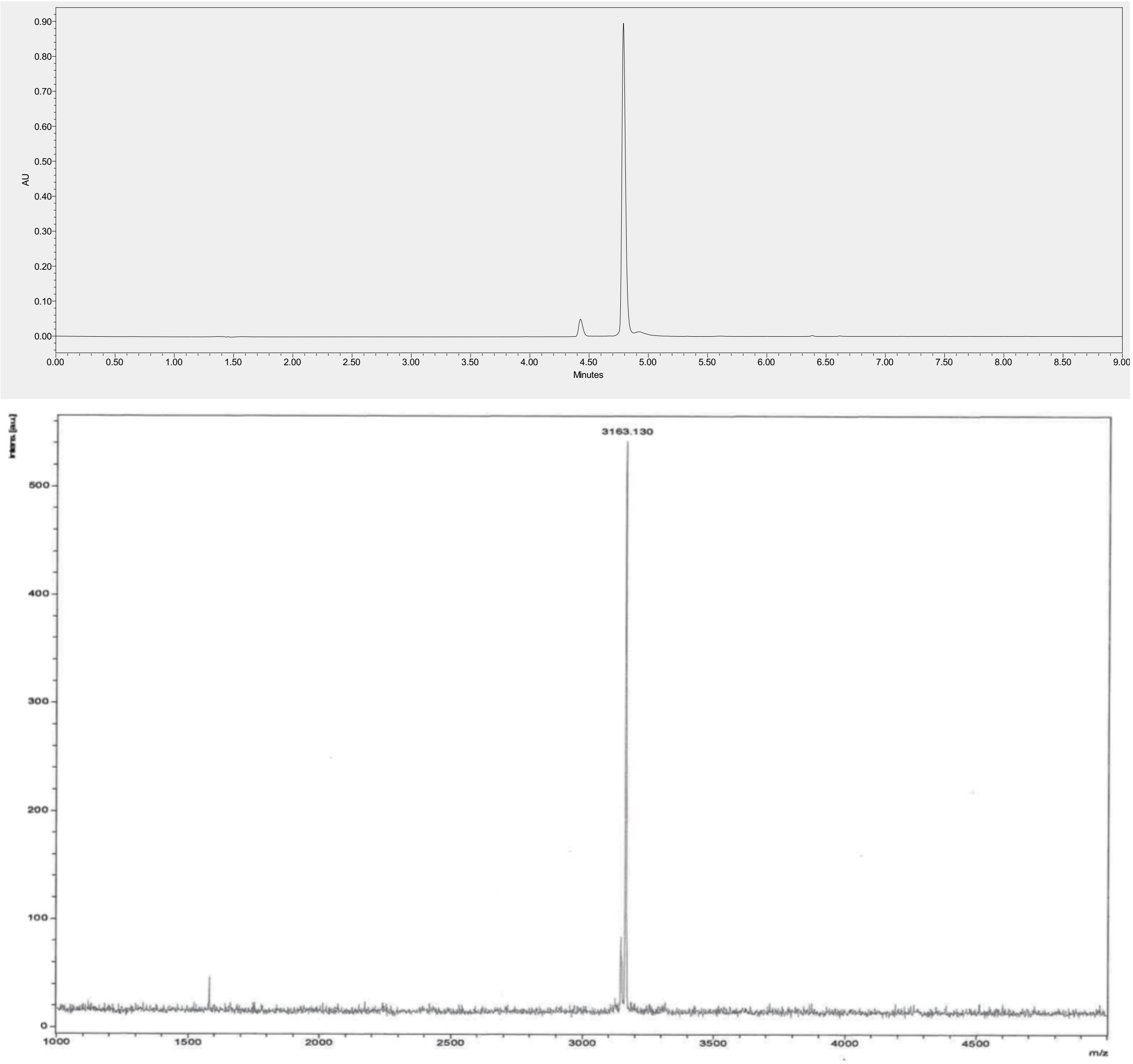


UPLC trace and MALDI-TOF MS for G1K-Q11E-V14L. Peptide had purity higher than 98%. The signal at lower m/z is impurity due to aspartimide formation.

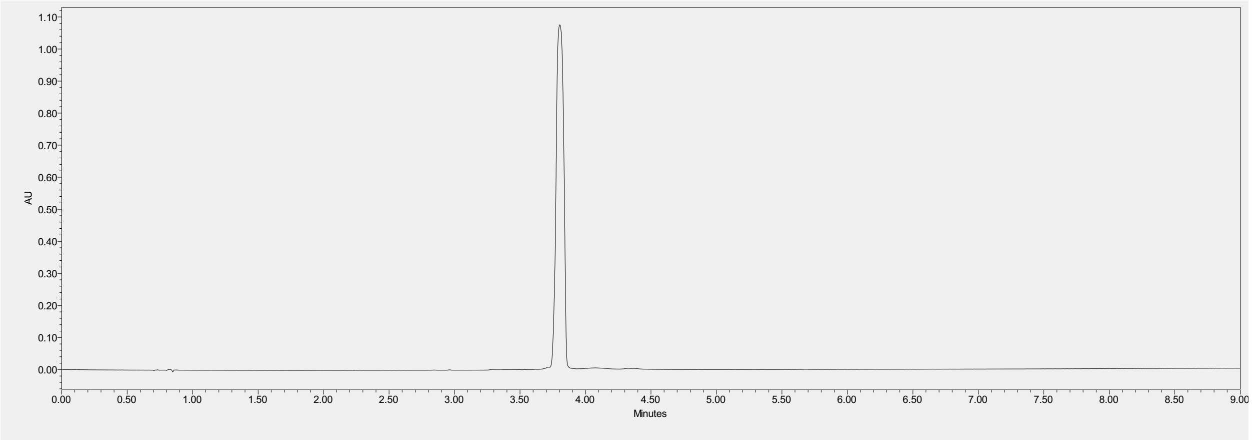


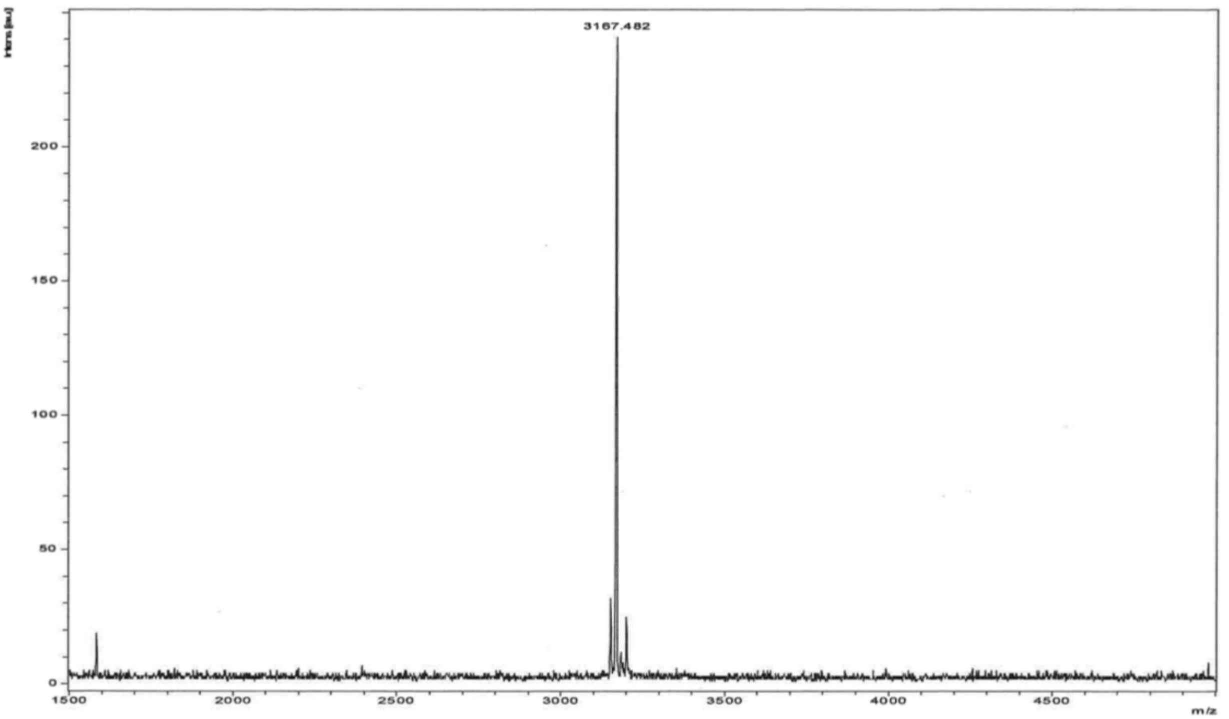


UPLC trace and MALDI-TOF MS for G1K-Q11E-V14I. Peptide had purity higher than 98%. The signal at lower m/z is impurity due to aspartimide formation.

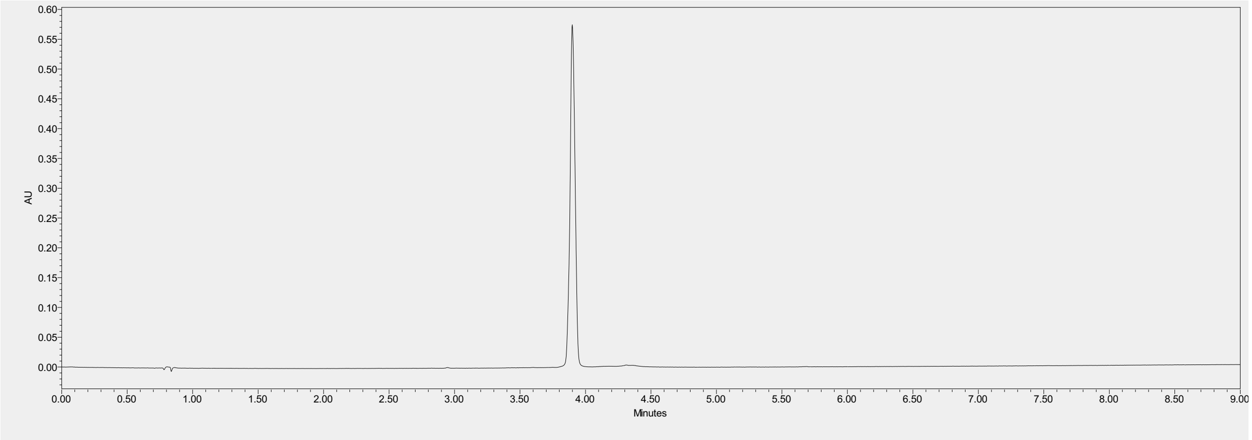


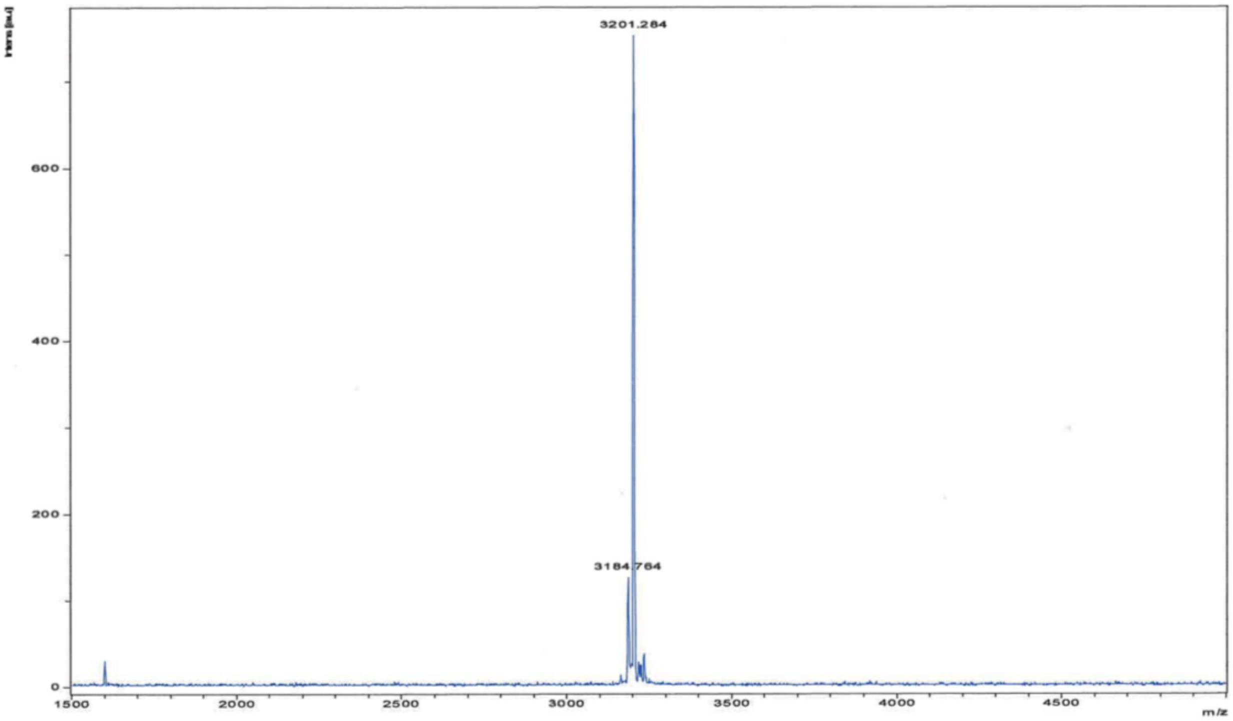


UPLC trace and MALDI-TOF MS for G1K-Q11E-V14F. Peptide had purity higher than 98%. The signal at lower m/z is impurity due to aspartimide formation.

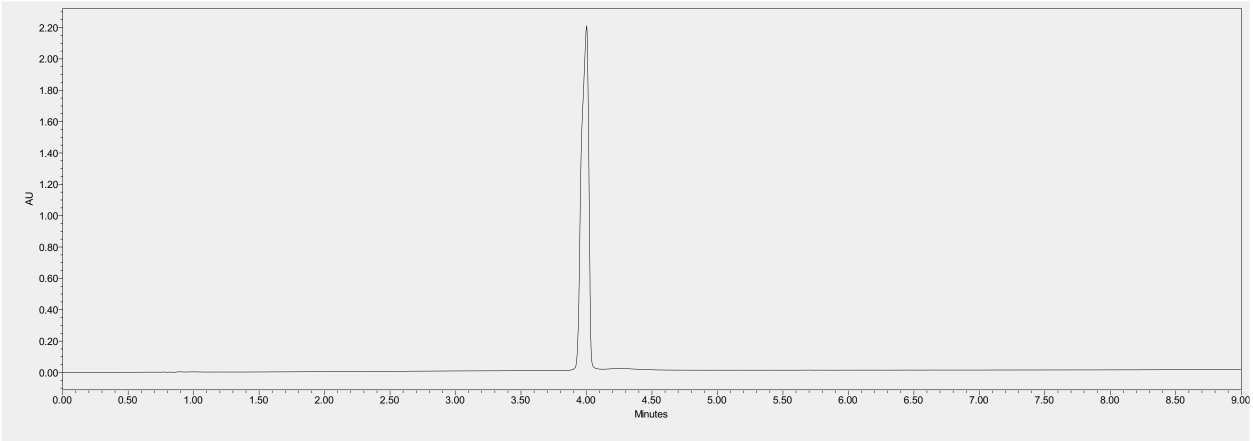


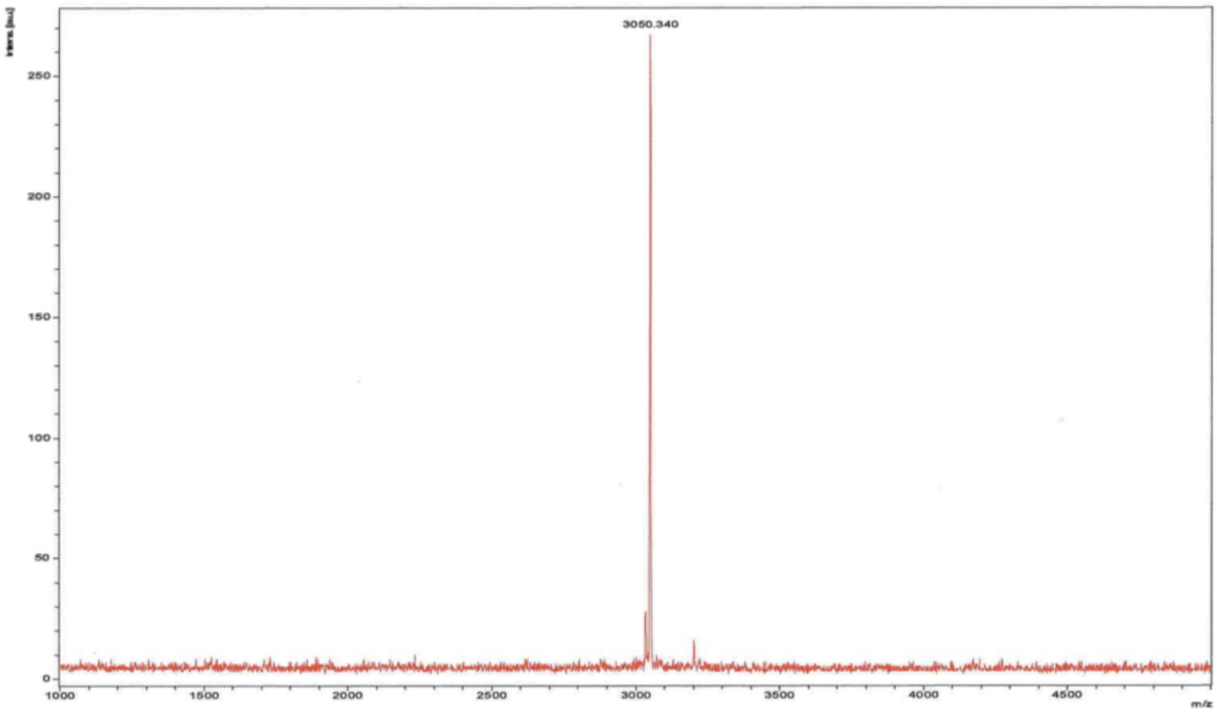


UPLC trace and MALDI-TOF MS for Q11E-V14A. Peptide had purity higher than 98%.

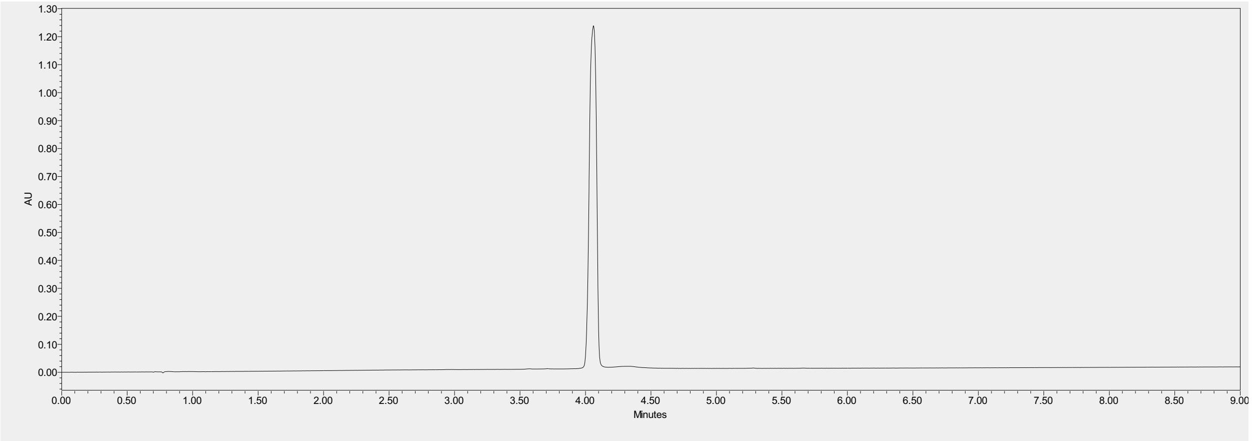


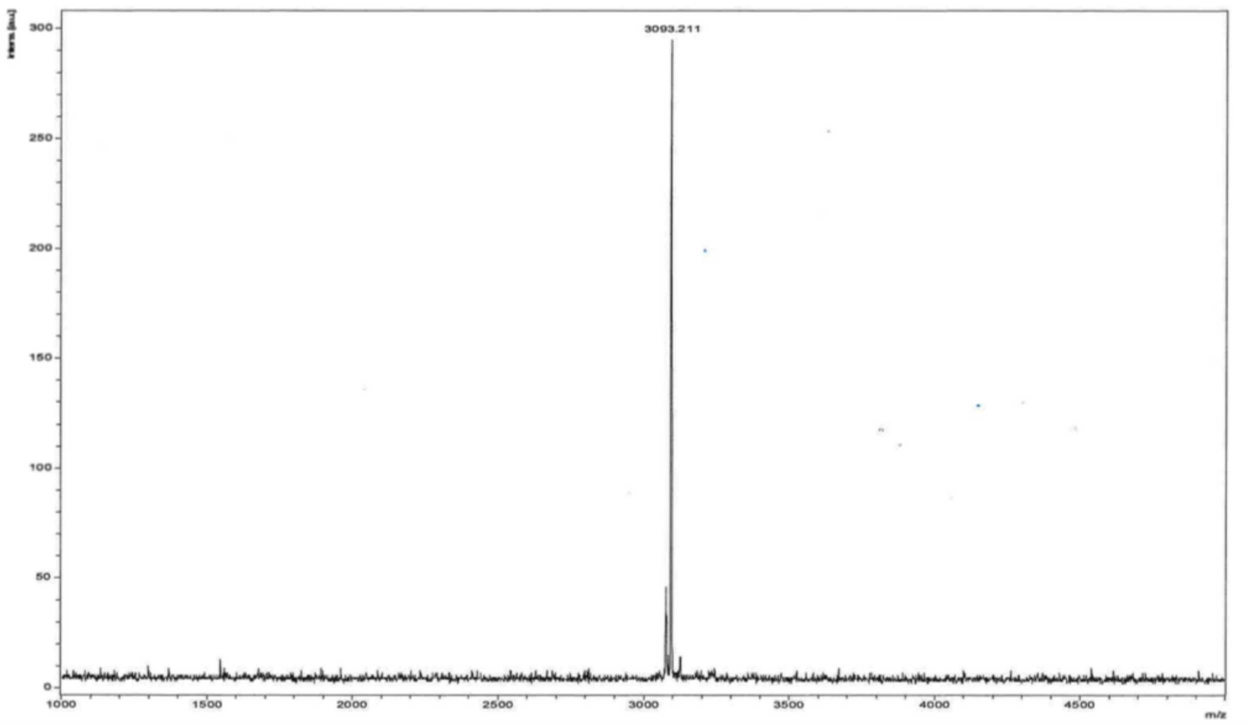


UPLC trace and MALDI-TOF MS for Q11E-V14L. Peptide had purity higher than 98%.
